# Early downregulation of hair cell (HC)-specific genes in the vestibular sensory epithelium during chronic ototoxicity

**DOI:** 10.1101/2025.05.05.652224

**Authors:** Mireia Borrajo, Erin A. Greguske, Alberto F. Maroto, Aïda Palou, Ana Renner, Víctor Giménez-Esbrí, David Sedano, Marta Gut, Anna Esteve-Codina, Beatriz Martín-Mur, Alejandro Barrallo-Gimeno, Jordi Llorens

## Abstract

Exposure of mammals to ototoxic compounds causes hair cell (HC) loss in the vestibular sensory epithelia of the inner ear. In chronic exposure models, this loss often occurs by extrusion of the HC from the sensory epithelium towards the luminal cavity. HC extrusion is preceded by several steps that begin with detachment and synaptic uncoupling of the cells from the afferent terminals of their postsynaptic vestibular ganglion neurons. The purpose of this study was to identify gene expression mechanisms that drive these responses to chronic ototoxic stress. We conducted four RNA-seq experiments that generated five comparisons of control versus treated animals. These involved two species (rat and mouse), two compounds (streptomycin and 3,3’-iminodipropionitrile, IDPN), and three time points in our rat/IDPN model. We compared differentially expressed genes and their associated Gene Ontology terms, and several genes of interest were validated by in-situ hybridisation and immunofluorescence analyses. Common and model-unique expression responses were identified. The earliest and most robust common response was downregulation of HC-specific genes, including stereocilium (*Atp2b2, Xirp2*), synaptic (*Nsg2*), and ion channel genes (*Kcnab1*, *Kcna10*), together with new potential biomarkers of HC stress (*Vsig10l2*). A second common response across species and compounds was the upregulation of the stress mediator *Atf3*. Model- or time-restricted responses included downregulation of cell-cell adhesion and mitochondrial ATP synthesis genes, and upregulation of the interferon response, unfolded protein response, and tRNA aminoacylation genes. The present results provide key information on the responses of the vestibular sensory epithelium to chronic ototoxic stress, potentially relevant to other types of chronic stress.

## INTRODUCTION

The vestibular system in the inner ear, also known as the labyrinth, is a sensor of head accelerations. The cells responsible for sensory transduction are known as hair cells (HC) due to the bundle of filiform extensions, named stereocilia, that are on the their apical end of the HC. Stereocilia bundles protrude into the endolymph-filled luminal cavities of the labyrinth and express the molecular mechanisms that trigger depolarization or hyperpolarization of the HC in response to their deflection caused by forces of bodily acceleration. In mammals, HCs are organized into five sensory epithelia per ear: one utricle, one saccule, and three cristae. There are two types of HC, type I (HCI) and type II (HCII), which can be differentiated by morphological, molecular, and physiological criteria (Eatock and Songer, 2011; Dalet et al., 2012; McInturff et al., 2018; Giunta et al., 2024). The most prominent differential feature is the calyx terminal, a unique afferent formation that encases HCIs but not HCIIs. These two types include several subtypes. For example, both HCIs and HCIIs may or may not express the calcium-binding protein oncomodulin (Hoffman et al., 2018) and calyces encasing HCIs may or may not express another calcium-binding protein, calretinin (Lysakowski et al., 2011).

Vestibular HCs and cochlear HCs are damaged by several toxic molecules, which are therefore named ototoxic compounds. These include aminoglycoside antibiotics (streptomycin, gentamicin, and others), the chemotherapeutic agent cisplatin, and several low molecular weight nitriles (3,3’-iminodipropionitrile - IDPN, *cis*-crotonitrile, allylnitrile) (Boyes et al., 2020). Human therapeutic use or animal experimental exposure to these ototoxic compounds cause HC loss resulting in permanent vestibular and auditory deficits, as HCs do not typically regenerate in mammals; if they do, it is only to a very limited, non-functional, extent. The molecular processes leading to the loss of HCs have been the subject of intense research, mostly in the hope that their understanding could reveal targets for prophylactic drugs that could prevent the permanent loss of function that may occur after the use of ototoxic agents. Most of the research has focused on ototoxic-induced HC apoptosis (Furness 2015). However, HCs can respond to injury stimuli by extrusion, a process by which the HC modifies its ultrastructural features and is ejected as a live cell from the sensory epithelium towards the vestibular luminal cavity. This is followed by the degeneration of the ejected cell in the lumen. Cell extrusion is a form of cell loss that can occur only in epithelia. It has been identified as the main process of the elimination of HCs in the inner ear of birds and amphibians (Hirose et al., 1999; Rubel et al., 2013) and has been occasionally observed in the inner ear of mammalian species after ototoxicity (Li et al., 1995; Nakagawa et al., 1997; Granados and Meza, 2005), but it has been scarcely studied. Recent evidence indicates that retinal photoreceptor cells may also suffer extrusion (Völkner et al., 2022). Previous research in our laboratory identified a model, the chronic exposure of rats to IDPN, in which most or all vestibular HCs are eliminated by extrusion (Seoane et al., 2001a,b). Subsequent research in rats (Sedó-Cabezón et al., 2015) and mice (Greguske et al., 2019) revealed that during IDPN intoxication the extrusion process is preceded by a reversible uncoupling of the HC and their post-synaptic afferents. A major hallmark of this process is the dismantling of the calyceal junction, a cell-adhesion and signalling complex that is normally formed between the HCI and its corresponding calyx. This is paralleled by a major loss of vestibular function (Sedó-Cabezón et al., 2015; Greguske et al., 2019; Martins-Lopes et al., 2019; Schenberg et al., 2023). If intoxication continues, stereocillia start to fuse and HCs are driven toward extrusion and irreversible loss (Llorens and Rodríguez-Farré, 1997; Seoane et al. 2001a, b). However, if intoxication is terminated before stereocilia fusion begins, then calyceal junction dismantlement and synaptic and behavioural defects are reversed (Sedó-Cabezón et al., 2015; Greguske et al., 2019; Martins-Lopes et al., 2019; Schenberg et al., 2023). Recently, we have collected evidence that calyx junction dismantlement also precedes HC extrusion in rats exposed to chronic streptomycin and may also occur in response to other sources of stress, such as stress caused in human vestibular epithelia by a neighbouring vestibular schwannoma (Maroto et al., 2023).

The observations above suggest that extrusion is an alternate process of regulated cell death similar to apoptosis. The genetically programmed process that executes apoptosis has been extensively characterized (Vitale et al., 2023), but the program driving HC extrusion has not yet been studied. Our observations that extrusion may be a slow process that begins with reversible steps make this study a particularly interesting one, as this opens doors for pharmacological intervention to stop damage progression or accelerate repair. The present study is a first approach to the discovery of the molecular mechanisms involved in vestibular sensory epithelium damage during chronic ototoxicity leading to HC extrusion. We hypothesized that these mechanisms would include specific gene expression signatures even though mechanisms other than gene expression modifications must be involved in the execution of HC extrusion. We also hypothesized that the most important changes in gene expression would be more conserved throughout evolution and more consistently triggered by different ototoxic compounds. Therefore, we performed RNAseq studies in chronic ototoxicity models involving two different species (mouse and rat) and two ototoxic compounds (IDPN and streptomycin). We also evaluated different time points over the course of the treatment in one of the models. As the objective was to understand the response to a slowly evolving stress condition, we reasoned that stress caused by the cell isolation protocols necessary for single-cell or purified cell population studies would mask the gene expression changes of interest. Therefore, we used whole tissue for the extraction and sequencing of RNA. The results show similar and dissimilar changes in gene expression in the different models. Among these, the earliest common response in all ototoxicity models is decreased expression of many HC-specific genes, and follow-up studies demonstrate decreased expression of their corresponding proteins. In addition, other common and model-unique responses to chronic ototoxicity were also identified.

## MATERIALS AND METHODS

### Animals and treatments

We used both Long-Evans rats, obtained from Janvier Labs (Le-Genest-Saint-Isle, France), and 129S1/SvImJ mice from a locally established colony, descendants of animals purchased from Jackson Laboratory (Bar Harbor, ME, USA). For *in vitro* experiments, neonatal rats were obtained from pairs of Long-Evans rats bred locally. In all experiments, the animals were housed in Macrolon cages in groups of at least 2 subjects with wood shavings as bedding. The housing conditions were standard, including a 12:12 h light:dark cycle (07:30-19:30 h), a room temperature of 22±2°C and *ad libitum* access to standard diet pellets (TEKLAD 2014, Harlan Laboratories, Sant Feliu de Codines, Spain).

Four RNA-seq experiments were carried out. In all of them, the animals were euthanized for tissue collection on the last day of treatment. We used exposure conditions that cause progressive vestibular damage as thoroughly described in previous publications (Sedó-Cabezón et al., 2015; Greguske et al., 2019; Maroto et al., 2023). In three experiments, we selected doses and exposure times that caused the most possible, but still reversible, loss of vestibular function after IDPN treatment in mice and rats, and after streptomycin treatment in rats. Thus, in the first experiment, 2-to 4-month-old male 129S1/SvImJ mice were given free access to water spiked with 0 (Control) or 30 (mouse/IDPN) mM IDPN (>98%, TCI Europe, Zwijndrecht, Belgium) as the sole source of drinking liquid for 8 weeks. In the second experiment, male adult Long-Evans rats (8-9 weeks old on reception) were given 0 (Control) or 20 (rat/IDPN-4wk) mM IDPN for 4 weeks. For the third experiment, male and female rats were obtained at post-natal day 8-9 and housed with their lactating mothers until weaning at day 21. Starting on day 21, the rats in the streptomycin group (rat/STREP) received streptomycin sulphate salt (Sigma-Aldrich, S6501), s.c., dissolved in phosphate-buffered saline (PBS), once a day for 6 weeks, at 500 mg/kg day (as free base). Control rats were injected with PBS. Initial injection volumes were 10 ml/kg to guarantee dose accuracy but were reduced to 2 ml/kg as the rats gained weight. The injection site in the dorsal skin was changed from day to day. Finally, the fourth rat/IDPN experiment was added to obtain data on an earlier step of HC detachment (Sedó-Cabezón et al., 2015) and for a later time point when stereocilia fusion indicates the beginning of the active HC extrusion (Seoane et al., 2001a; Sedó-Cabezón et al., 2015). In this fourth experiment, the 20 mM concentration was used for 3 weeks (rat/IDPN-3wk) or 7 weeks (rat/IDPN-7wk) and a single control group included animals euthanized at each time point.

### Assessment of vestibular function

To evaluate vestibular function in rats, we assessed the tail-lift reflex. Briefly, rats are held by the base of the tail and quickly but gently lifted upward to approximately 40 cm above ground and then immediately returned down. In healthy rats, this triggers a trunk and limb extension reflex as a landing response. Loss of vestibular function results in the loss of this anti-gravity extension reflex, and the rat curls ventrally. The reflex response is recorded with high-speed video and the movie is used to determine the tail-lift angle (TLA). The TLA is determined by the minimum angle drawn from the nose, to the back of the neck and the base of the tail during the tail-lift manoeuvre, as described (Martins-Lopes et al., 2019; Maroto et al., 2021 a,c).

In mice, loss of vestibular function was assessed using a previously validated behavioural test battery (Soler-Martín et al., 2007; Saldaña-Ruíz et al., 2013; Greguske et al., 2019; Maroto et al., 2021c). Briefly, mice are blindly rated from 0 to 4 on 6 items. Ratings are scaled as: 0 for the behaviours that characterize normal healthy animals and 4 for the maximum possible deviation from normal behaviour. The mice are placed for one minute in a clean open arena, and rated for circling (stereotyped circulatory ambulation), retropulsion (backward movement), and abnormal head movements (intermittent extreme backward extension of the neck). Afterwards, the animals are rated for the tail-lift reflex, the contact inhibition of the righting reflex, and the air-righting reflex. More detailed descriptions of the battery are available (Saldaña-Ruiz et al., 2013; Greguske et al., 2019; Maroto et al., 2021c). Although the tail-lift reflex included in the mouse test battery is the same reflex used to assess vestibular dysfunction in rats, a protocol for reliable quantification of this reflex has not been validated in mice yet.

### Sample collection

At the end of the exposure periods, animals were decapitated and their vestibular epithelia were quickly dissected in ice-cold PBS for RNA isolation. From each animal, the six cristae and the two utricles were isolated, cleaned of adjacent tissues, pooled, placed in an Eppendorf tube, frozen in liquid nitrogen, and stored at −80° C. After RNA isolation and quality control as described below, three samples per group were used for RNA-seq. Thus, mouse/IDPN, rat/IDPN-4wk and rat/STREP experiments compared 3 control and 3 treated animals. In the streptomycin experiment, both the control and treated groups were composed of two male and one female rat. The other rat/IDPN experiment compared 3 control rats, 3 rats exposed to IDPN for 3 weeks, and 3 rats exposed to IDPN for 7 weeks.

For immunofluorescence and RNA-scope studies, vestibular epithelia were quickly dissected in cold 4% paraformaldehyde in PBS under a fume hood, fixed in the same solution for 1 h, transferred into a cryo-preservative solution (34.5% glycerol, 30% ethylene glycol, 20% PBS, 15.5% distilled water) and stored at −20°C. The present experiments were carried out using new samples as well as archival samples from previous studies (Sedó-Cabezón et al., 2015; Greguske et al., 2019; Maroto et al., 2023) stored for up to 10 years. For each analysis, we used sensory epithelia from control and treated animals that had been obtained at the same time.

### Short read, low-input RNA sequencing

Total RNA was extracted from the samples using the Qiagen RNeasy mini kit and the manufacturer’s protocol. RNA sequencing libraries were prepared following the SMARTseq2 protocol (Picelli et al., 2013) with some modifications. Briefly, total RNA samples were quantified by the Qubit® RNA BR Assay kit (Thermo Fisher Scientific) and RNA integrity was estimated with an Agilent RNA 6000 Pico Bioanalyzer 2100 Assay (Agilent). Reverse transcription of the total RNA input material of 1.8 µl (6-11 ng, depending on sample availability) was performed using SuperScrpit II (Invitrogen) in the presence of oligo-dT30VN (1µM; 5′-AAG CAG TGG TAT CAA CGC AGA GTA CT30VN-3′), template-switching oligonucleotides (1 µM) and betaine (1M). The cDNA was amplified using the KAPA Hifi Hotstart ReadyMix (2x) (Roche) and 100 nM IS PCR primer (5′-AAG CAG TGG TAT CAA CGC AGA GT-3′) with 8 cycles of PCR amplification. Following purification with Agencourt Ampure XP beads (1:1 ratio; Beckmann Coulter), the product size distribution and the quantity were assessed on a Bioanalyzer High Sensitivity DNA Kit (Agilent). The amplified cDNA (200 ng) was fragmented for 10 min at 55 ° C using Nextera XT (Illumina) and amplified for 12 cycles with indexed Nextera PCR primers. The library was purified twice with Agencourt Ampure XP beads (0.8:1 ratio) and quantified on a Bioanalyzer using a High Sensitivity DNA Kit.

The libraries were sequenced on the HiSeq 4000 or HiSeq 2500 (Illumina) in paired-end mode. The read length for the HiSeq 4000 runs was 2×76 bp, using the HiSeq 4000 SBS Kit and HiSeq 4000 PE Cluster Kit sequencing flow cell. When using the HiSeq 2500, the libraries were sequenced in paired-end mode with a read length of 2×76 bp using the TruSeq SBS Kit v4.

Primary data analysis, including image analysis, base calling and quality scoring, was performed using the manufacturer’s software: Real-Time Analysis (RTA 2.7.7) for HiSeq 4000 and RTA 1.18.66.3 for HiSeq 2500. This was followed by the generation of FASTQ sequence files.

### RNA-seq data processing and analysis

Mouse RNA-seq reads were mapped against the Mus musculus reference genome (GRCm38) with STAR/2.5.3a (Dobin et al., 2013) using ENCODE parameters. Genes and isoforms were quantified with RSEM/2.3.0 (Li and Dewey, 2011) with default parameters using the gencode.M15 annotation. Likewise, rat RNA-seq reads were mapped against the Rattus norvegicus reference genome (Rnor6.0) while the Rnor6.0.92 annotation was used for gene and isoform quantification. Original raw and processed data are available from Gene Expression Omnibus, accession number GSE292474 (https://www.ncbi.nlm.nih.gov/geo/query/acc.cgi?acc=GSE292474)

Differential expression analysis was performed with the R package DESeq2/1.18 (Love et al., 2014). The regularized log transformation of the counts was used for plotting. Genes with FDR<5% and |FC|>1.5 were considered significantly differentially expressed. Heatmaps were drawn with the R package ‘ggplot2’ using Z-score normalisation and the PCA was done with the ‘prcomp’ R function. Gene ontology (GO) enrichment analyses of the differentially expressed genes were performed with g:Profiler (Raudvere et al., 2019). We separately analysed the lists of upregulated or downregulated genes from each control vs treated comparison. Enrichment analyses were done also on the lists of genes similarly or differentially regulated across experiments (for instance, genes downregulated in both the streptomycin and the IDPN-4wk groups). To reduce redundancy in the lists of GO terms, the REVIGO tool (Supek et al., 2011) was used to cluster groups by semantic similarity analysis and thus generate shorter lists of cluster representatives.

In addition to GO analyses, Gene Set Enrichment Analyses (GSEA) (Subramanian et al., 2005; Mootha et al., 2003) were performed with the original expression datasets to evaluate whether the gene expression changes induced by the treatments matched any well-known biological process as defined by the Hallmark Gene Sets.

### Vestibular epithelium culture

Vestibular epithelia were cultured as described previously (Borrajo et al., 2024). The epithelia were dissected from rats on post-natal day 1 in Leibovitz’s L-15 medium (Gibco, cat.# 11415064) and cultured free-floating in a standard culture medium composed of Dulbecco’s Modified Eagle’s Medium-Nutrient Mixture F12 (DMEM/F12) (Gibco, cat. #11320033) supplemented with 25 mM HEPES (Gibco, cat.# 15630080), 2% N2 supplement (Cell Therapy Systems, cat.# A1370701), 1% GlutaMAX (Gibco, cat.# 35050061), 2 g/L glucose (Sigma, cat.# 49163) and 1.5 g/L penicillin (ThermoFisher, cat.#J63032.14), in a 5% CO2/95% air environment at 37°C. This medium was enriched only for the four initial days with 50 ng/mL EGF (Preprotech, cat.# AF100-15), 0.1 μg/mL FGF2 (Sigma, cat.#F0291) and 0.1 μg/mL FGF10 (Preprotech, cat.# AF-100-26), as well as with 1 mMN-acetyl-L-cistein (Sigma, cat.#A9165), 10mM nicotinamide (Sigma, cat.# N0636) and 1% insulin-transferrin-selenium-ethanolamine (ITS-X) supplement (Gibco, cat.# 51500056). The cultures were exposed to 0 or 0.05 μM streptomycin (from streptomycin sulphate salt, Sigma-Aldrich, cat.# S6501) from days 14 to 21 *in vitro*. This concentration of streptomycin was selected as the maximal concentration causing no loss of HCs, according to unpublished data. At the end of the treatment, tissues were fixed and stored for RNA-scope as described above for samples from the *in vivo* studies.

### Immunohistochemical labelling

For immunohistochemistry of the vestibular sensory epithelia, the following commercial primary antibodies were used at the indicated dilutions: mouse monoclonal anti-contactin-associated protein (CASPR1) (clone K65/35, cat.#MABN69, Millipore, RRID: AB_2083496, 1/400), mouse monoclonal anti-myosin-7A (MYO7A) (cat.# 138-1, supernatant, Developmental Studies Hybridoma Bank, RRID: AB_2282417, 1/100), rabbit anti-potassium voltage-gated channel subfamily A member 10 (KCNA10, Kv1.8)(cat.# APC-157, Alomone Labs, RRID: AB_2341039, 1/200), rabbit anti-MYO7A (cat.# 25-6790, Proteus Biosciences, RRID: AB_10015251, 1/400), rabbit anti-oncomodulin (cat.# OMG4, Swant, RRID:AB_10000346, 1/400), rabbit anti-plasma membrane calcium transporting ATPase 2 (PMCA2, encoded by the Atp2b2 gene) (cat.# PA1-915, Invitrogen, RRID: AB_2243199, 1/400), goat anti-Delta/Notch Like EGF Repeat Containing (DNER) (cat.# AF2254, R&D Systems, RRID:AB_355202, 1/200), goat anti-Osteopontin (encoded by the Spp1 gene) (cat.# AF808, R&D Systems, RRID: AB_2194992, 1/200), and guinea pig anti-calretinin (cat.# 214104, Synaptic Systems, RRID: AB_10635160, 1/500). In addition, a rabbit anti-VSIG10L2 antibody was generated against the VSXL-756:772 sequence (DPIQESIDAPVNVTITV) using the ThermoFisher Custom Antibodies services. To reveal the primary antibodies, we used appropriate fluorochrome-conjugated donkey secondary antibodies from Invitrogen / Thermo Fisher. These included DayLight-405, and Alexa-Fluor −488, −555, and −647 fluorochromes for four-channel imaging.

The entire vestibular epithelia were immunolabelled as previously described (Greguske et al., 2019; Maroto et al., 2021, 2023). To avoid biases from batch-to-batch differences in the label, equal numbers of specimens from each experimental group were processed in parallel. Briefly, the main protocol included permeation and blocking in 4% Triton-X-100 (Lysakowski et al., 2011) and 20% donkey serum in PBS for 90 min at room temperature, followed by incubation with the mixed primary antibodies in 0.1% Triton-X-100 and 1% donkey serum in PBS for 48 h at 4°C, then by incubation with the secondary antibodies in 0.1% Triton-X-100 in PBS overnight at 4°C. Specimens were thoroughly rinsed in PBS following each incubation and gently rocked during all incubations and rinses. To assess KCNA10 and PCMA2 immunoreactivity, the samples were first permeated at room temperature with 4% Triton-X100 for 1 h, then blocked with 1% fish gelatin (cat.# G7765, Sigma-Aldrich) for 1 h (Lysakowski et al., 2011). In this protocol, primary and secondary antibody solutions contained 0.1% Triton-X-100 and 1% fish gelatin. After immunolabelling, some batches of samples were whole-mounted in Fluoromount medium (F4680, Sigma-Aldrich), while other samples were sectioned before mounting. For sectioning, specimens were placed in 0.49% gelatin, 30% bovine serum albumin, and 20% sucrose in PBS overnight at 4°C. The next day, the sample was oriented and included in between two fused blocks formed by solidification of the same mixture with 2 % glutaraldehyde. The block was then used to obtain 40 μm-thick sections in a Leica VT1000S vibrating microtome. This protocol has been described in detail elsewhere (Maroto et al., 2021b).

### RNA scope in-situ hybridization

Vestibular epithelia from *in vivo* or *in vitro* studies were processed for simultaneous immunofluorescence and RNAscope in-situ hybridization. We used the RNAscope Multiplex Fluorescent Reagent Kit v2 (cat. # 323100, ACD Bio-Techne) with the following probes: RNAscope® Probe - Rn-Vsig10l2-C1,PREDICTED: Rattus norvegicus V-set and immunoglobulin domain containing 10 like 2 (Vsig10l2) mRNA (cat. # 1241271-C1), RNAscope® Probe-Rn-Atf3 (cat. #414981) and Rn-Ppib-C3 - Rattus norvegicus peptidylprolyl isomerase B (Ppib) mRNA (cat. # 313921-C3) or RNAscope® Positive Control Probe - Rn-Ppib-C2 - Rattus norvegicus peptidylprolyl isomerase B (Ppib) mRNA (cat. #313921-C2) as a positive control. To label the probes, we used the fluorophores Opal 520 (cat. # FP1487001KT, Akoya Biosciences), Opal 570 (cat. # FP1488001KT) and Opal 690 (cat. # FP1497001KT). Briefly, free-floating epithelia were washed in RNAscope Wash Buffer and dehydrated with graded concentrations of ethanol. The samples were then washed and incubated in H_2_O_2_ 0.3% (v/v) for 10 min at RT. Sections were then hybridized in probe mixture (either Rn-Vsig10l2-C1 plus Rn-Ppib-C3 or Rn-Atf3 plus Rn-Ppib-C2 at a 50:1 concentration, as indicated by the manufacturer) inside screw microtubes for 2 h at 37°C using a water bath. Afterwards, tissues were washed and incubated in amplification solutions Amp1, Amp2, and Amp3 for 35, 20, and 35 min, respectively, at 37°C. For signal detection, HRP-C1 and HRP-C3/HRP-C2 solutions were added to the sections for 15 min each at 37°C, followed by a 30 min incubation of TSA-diluted Opal 520 and Opal 570/Opal 690 at 37°C. After washing, the sections were processed for MYO7A immunostaining with the rabbit MYO7A antibody and mounted following the protocol described above.

### Confocal microscopy imaging and analysis

Vestibular epithelia were observed in a Zeiss LSM880 spectral confocal microscope. The 63x (Numeric Aperture =1.4) objective was used to obtain images of 135 x 135 μm or 67.5 x 67.5 μm (zoom 2). Z-stacks of optical sections were obtained at 0.35 μm interval spanning 6 to 30 μm as required for the planned analysis. The same image acquisition settings were used for all samples from each immunolabelling batch. To measure fluorescence intensities, we used ImageJ software (National Institute of Mental Health, Bethesda, Maryland, USA). The expression of the plasma membrane calcium-transporting ATPase (PMCA2), encoded by the *Atp2b2* gene, was evaluated in whole-mount utricles (one utricle per animal) labelled with mouse anti-MYO7A, anti-calretinin and anti-PMCA2 antibodies. From each utricle 15 different areas were acquired with Z-stacks of 30 optical sections. These were then used to generate maximal intensity projections of 14-section sub-stacks for quantitative analysis. In each image, a rectangular region of interest (ROI) of 12.6 x 7.8 μm was used to assess the fluorescence intensity of the PMCA2 channel in a stereocilia bundle and the background fluorescence intensity was subtracted. From each image, at least 12 different bundles were quantified, so PMCA2-associated fluorescence for each utricle was estimated as the mean value of at least 180 (usually more than 200) individual hair bundles. To evaluate the effects of the treatments, we compared 3 control rats with 4 IDPN-4wk rats and 4 control rats with 4 streptomycin rats.

DNER expression was quantified using whole-mount cristae labelled with rabbit anti-MYO7A, anti-CASPR1, anti-calretinin and anti-DNER antibodies. From each crista, four images were acquired, two from the central part and two from the periphery, with Z-stacks extending up to 30 μm. For quantitative analysis, sub-stacks of 6 sections were made to select the level of the epithelium containing the supra-nuclear region of HC, and an oval ROI (diameters of 2,7 and 2,1 μm) was used to quantify fluorescence intensity of at least 60 HC per image. To evaluate the effects of the treatments, we compared 3 control rats with 4 IDPN-4wk rats, and 4 control rats with 4 streptomycin rats.

KCNA10 expression was assessed in whole-mount cristae labelled with anti-osteopontin, anti-CASPR1, and anti-KCNA10 antibodies. From each crista, four images were obtained as described above for DNER. For quantitative analysis, sub-stacks of 7 sections were made to select the level of the epithelium containing the mid-lower nuclear region of HCI, to obtain circular slices of the calyceal junction adhering HCI with the afferent calyx. The ROI used for fluorescence measurement was made up of three circles, 0.60 μm in diameter each, placed at the vertices of an equilateral triangle with 4.1 μm edges. From each cell, two measurements were obtained, so the estimation of the fluorescence intensity was derived from 6 readings, and (approximately) evenly placed around the cell’s basolateral membrane. From each image, a mean fluorescence intensity was obtained by averaging the readings of 100 HCI. To evaluate the effects of the treatments, we compared 3 control rats with 3 IDPN-4wk rats and 3 control rats with 3 streptomycin rats.

For MYO7A, osteopontin and calretinin expression analyses, 40 μm-thick sections of crista samples were imaged and fluorescence intensities determined as described in detail elsewhere (Maroto et al., 2023). Briefly, 3 μm thick Z sub-stacks were used to assess the fluorescence intensity of 30 HCs per rat with a rectangular ROI. Data were normalized to control mean values of the same labelled batch to circumvent batch-to-batch differences in immunostaining. These analyses included 5 to 8 control rats and 6 to 10 IDPN-4wk rats.

The expression of the VSIG10L2 protein was evaluated in utricles (9 control and 9 IDPN-4wk samples) labelled with anti-VSIG10L2, anti-osteopontin, anti-calretinin and monoclonal anti-MYO7A antibodies. For quantitative analysis, 27 HCs were selected from each image using a pre-defined grid. For each cell, the fluorescence intensity above background was measured in the supranuclear zone using a circular ROI (5 μm in diameter).

For RNAscope analysis, Z-stacks of 50 optical sections were taken, obtained every 0.35 μm. For each epithelium, 4 images were collected in total: 2 from the centre of the epithelium and 2 from the peripheral zone. Then, the HC spanning section was selected by creating maximum intensity projections. For ATF3 expression analysis, the number of PPIB and ATF3 puncta was measured using the “Find maxima” tool of ImageJ software, after thresholding for intensity. The total number of puncta was averaged by individual, area, and number of HCs. To quantify VSIG and PPIB fluorescence intensity, images were thresholded to select VSIG or PPIB puncta. Then the mean grey value (that is, the fluorescence intensity) was acquired using the “measure” function of ImageJ. This number was averaged by individual, area, and total cell count.

### Evaluation of HC loss

To estimate the amount of HC loss that occurs in the mouse/IDPN exposure condition, we counted the number of hair bundles visible in scanning electron microscopy (SEM) images at 1500X magnification from the central area of the utricle of mice from previously published experiments (Greguske et al., 2019). For the rat/IDPN-4wk condition, HC counts were obtained in the cristae and utricles of an independent batch of control and treated rats, immunostained with anti-MYO7A and anti-calretinin antibodies and imaged at the confocal microscope as described above. Five 67.5 x 67.5 μm images per animal were collected and counted: central crista, peripheral crista, utricle striola, and lateral and medial peripheries of the utricle. For the rat/STREP condition, the samples used for counting were from rats treated together with the animals used for the RNA-seq analysis. They were processed as described for the rat/IDPN-4wk condition, except that a single image from the peripheral region of the utricle was obtained.

### Statistics

Padj / FDR values were used to evaluate gene and GO enrichment. For other data, such as immunofluorescence intensity data, group comparisons were done using Student’s t-test, corrected for multiple comparisons if needed, or ANOVA test as appropriate, using the GraphPad Prism software. In all cases, the significance level was set at p<0.05.

## 2. RESULTS

### 2.1. SYSTEMIC AND VESTIBULAR TOXICITY

#### Subchronic ototoxic exposure caused a progressive loss of vestibular function with limited loss of HCs and limited systemic toxicity

In four separate experiments, we obtained RNAseq data for five different treated *vs.* control comparisons. These included 3 different models of subchronic ototoxicity (mouse/IDPN, rat/IDPN, and rat/STREP) and 3 different time points (3, 4, and 7 weeks) of the rat/IDPN model. The mouse/IDPN, rat/IDPN-4wk, and rat/STREP experiments were performed to investigate the gene expression changes associated with the behavioural and histological effects described in previous studies (Sedó-Cabezón et al., 2015; Greguske et al., 2019; Maroto et al., 2023). The rat/IDPN-3wk and rat/IDPN-7wk groups were studied as models of earlier and later stages of the chronic ototoxicity process, respectively. As described previously (Sedó-Cabezón et al., 2015; Greguske et al., 2019; Martins-Lopes et al., 2019; Maroto et al., 2023), the exposed animals did not show overt signs of systemic toxicity, except a reduced increase in body weight compared to controls in some cases (Suppl. Fig. S1). Also, as revealed in mice by their vestibular dysfunction ratings (Saldaña-Ruiz et al., 2013) and in rats by the tail-lift reflex (Martins-Lopes et al., 2019; Maroto et al., 2021), the treated animals showed the expected progressive decline in vestibular function (Fig. 1 A-D).

**Figure 1.**
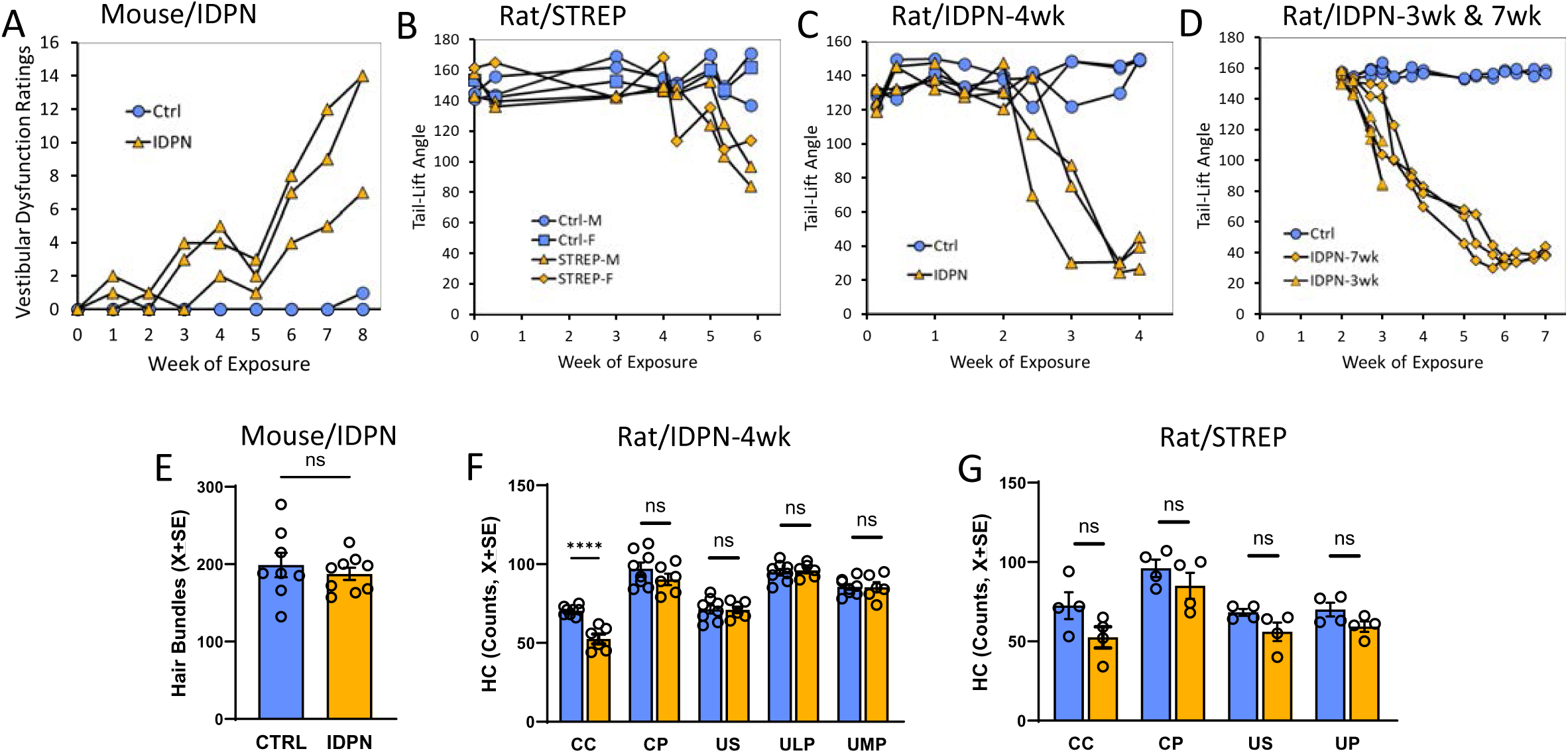
Effects of subchronic ototoxicity on vestibular function and HC density in the vestibular sensory epithelium. **A-D:** Vestibular function; graphs show the Vestibular Dysfunction Ratings (in mice, **A**) or the Tail-Lift Angle (in rats, **B**, **C**, and **D**) of the individual animals from which the RNA was extracted for RNA-seq analysis. **E:** Number of hair bundles counted in 1500X magnification SEM images of the central region of the utricles of mice exposed to 0 (CTRL) or 30 mM IDPN. **F:** Number of HCs (MYO7A+) in different regions of the vestibular epithelia from rats exposed to 0 or 20 mM of IDPN for 4 weeks. CC: crista centre, CP: crista periphery, US: utricle striola, ULP: utricle lateral periphery, UMP: utricle medial periphery. **G:** Number of HCs (MYO7A+) in different regions of the vestibular epithelia from rats exposed to 0 or 500 mg/kg/day of streptomycin for 6 weeks. CC: crista centre, CP: crista periphery, US: utricle striola, UP: utricle periphery.

Except for the rat/IDPN-7wk group, selected dosing schedules were expected to cause no or only limited loss of HCs, according to previous data (Sedó-Cabezón et al., 2015; Martins-Lopes et al., Greguske et al., 2019; Maroto et al., 2023). To corroborate this assertion, we collected additional data as shown in Fig. 1 E-G. For the mouse/IDPN experiment, only a rough estimate based on SEM images from our previous study (Greguske et al., 2019) was available. The number of hair bundles in the central region of the utricle was not modified after 8 weeks of exposure (Fig. 1E, t=0.670, 15 df, p=0.513). However, a significant presence of extruding profiles was observed. These extruding HCs were likely determined to be eliminated, because a significant 22% reduction in hair bundle numbers was recorded in utricles of mice that were evaluated 13 weeks after the end of the treatment (154.3±6.0 *vs*. 198.9±15.8 in controls, t=2.32, 12 df, p=0.038). In rats exposed to IDPN, detailed numbers of HCs were obtained after 4 weeks of exposure. As shown in Fig. 1F, a significant decrease in HC counts was recorded in the central region of the crista only (25%, t=6.17, 12 df, p<0.0001), but not in the other regions (all t’s < 1.20, 12 df, all p’s>0.254). In this case, a negligeable proportion of HCs were determined to be eliminated, because when IDPN-4wk rats were allowed to survive for 7 weeks after treatment, all HC counts were equal to those obtained at the end of the 4-week exposure period, with a significant decrease in the central region of the crista only (54.8+1.76 vs. 70.4+1.03 in controls, t=7.15, 16 df, p<0.0001). After streptomycin exposure (Fig. 1G), HC counts did not show significant differences in any of the regions of the vestibule (all t‘s < 1.98, 6 df, all p’s > 0.095), but the mean counts of the treated rats were 11 to 18% lower than the mean counts of control rats.

### 2.2. RNA-seq: OVERALL ANALYSIS

#### RNA-seq analyses identified gene sets with altered expression after subchronic ototoxic exposure

In all four experiments, treated animals showed significant changes in gene expression relative to control animals. In every case, principal component analysis (PCA) using the 500 most variable genes (Fig. 2A-D) resulted in a first component (PC1) that collected more than half of the variance (range 52.2 to 80.3 %) and ordered control and treated animals (n=3 in all groups). Heat maps of the 50 most significantly altered genes in each comparison between control and treated animals successfully grouped the samples according to treatment group (Fig. 2E-I). For each of these comparisons, the total number of differentially expressed genes (DEGs) and the numbers of downregulated and upregulated genes are shown in Table 1. Volcano plots, shown in Fig. 3, display these DEGs according to log2-Fold_Change and log-Padj. Comparisons across experiments revealed the presence of a significant batch effect, so the raw data could not be merged for a global analysis. Therefore, GO analyses with g:Profiler were performed separately for each treated *vs.* control comparison. Afterwards, comparisons across experiments were done by comparing both the lists of DEGs and the results of the GO analysis from each experimental model. Lists of DEGs and GO terms shared or differing across experiments or groups of experiments were obtained and evaluated as detailed throughout the text.

**Figure 2.**
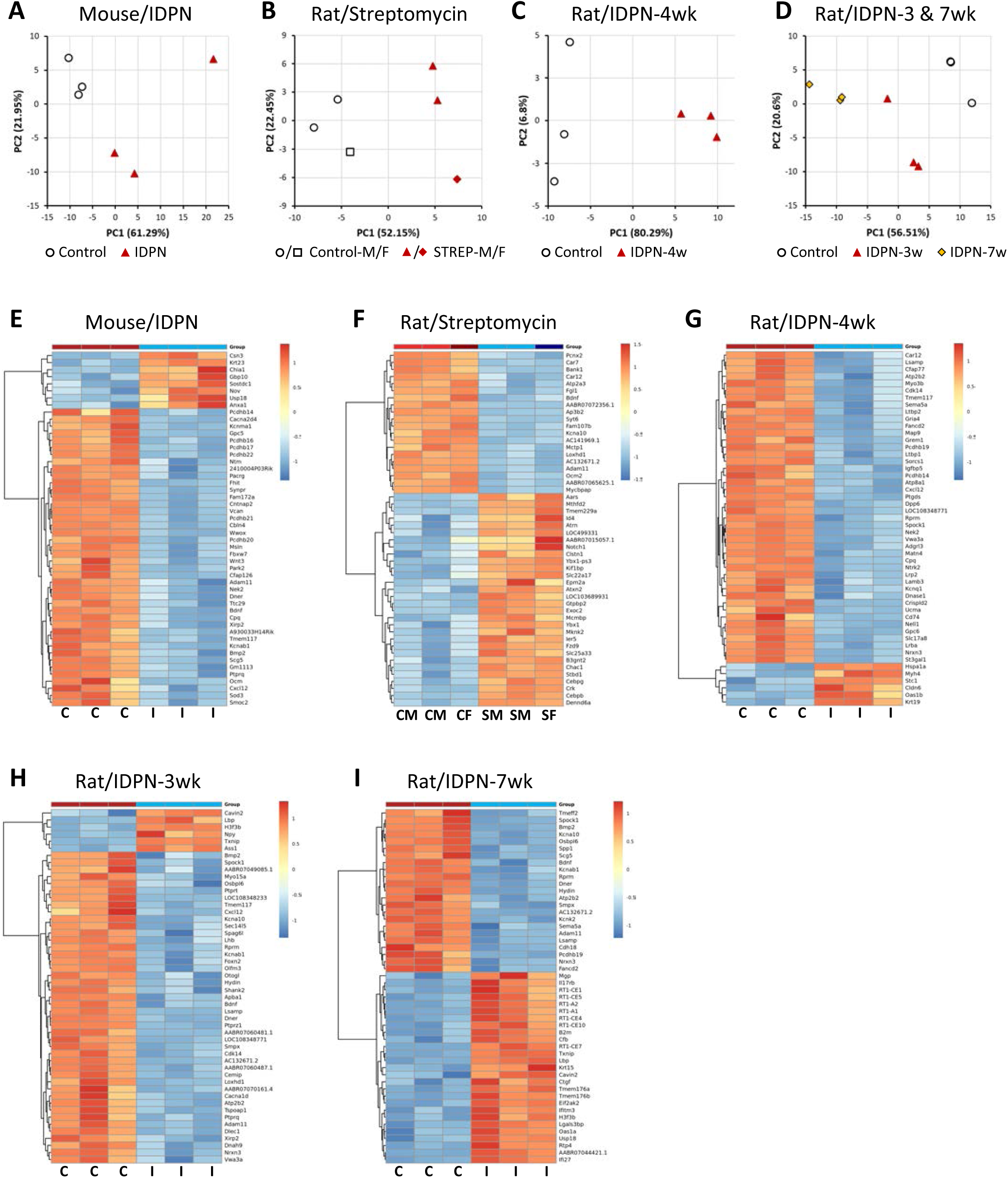
Effects of chronic ototoxicity on gene expression in vestibular sensory epithelium determined by RNA-seq analysis of control and treated rats or mice (n=3/group). **(A-D):** Principal Component Analysis (PCA) with the 500 most variable genes in four individual experiments, mice treated with IDPN **(A)**, rats treated with streptomycin **(B)**, rats treated with IDPN for 4 weeks **(C)** and rats treated with IDPN for 3 or 7 weeks **(D)**. Control and treated animals distributed according to PC1. The percentage figures are the percent of the total variance accounted for by the component. In the streptomycin experiment, M and F identify male and female rats, respectively. **(E-I):** Expression Heat Maps of the 50 most significantly altered genes in the five control vs treated comparisons emerging from the four experiments in **A-D**. Thus, data are shown comparing IDPN mice **(E)**, streptomycin rats **(F)**, rats treated with IDPN for 4 weeks **(G)**, 3 weeks **(H)**, or 7 weeks **(I)** with control animals in the experiment. C: control, I: IDPN, S: streptomycin, M: male, F: female.

**Figure 3.**
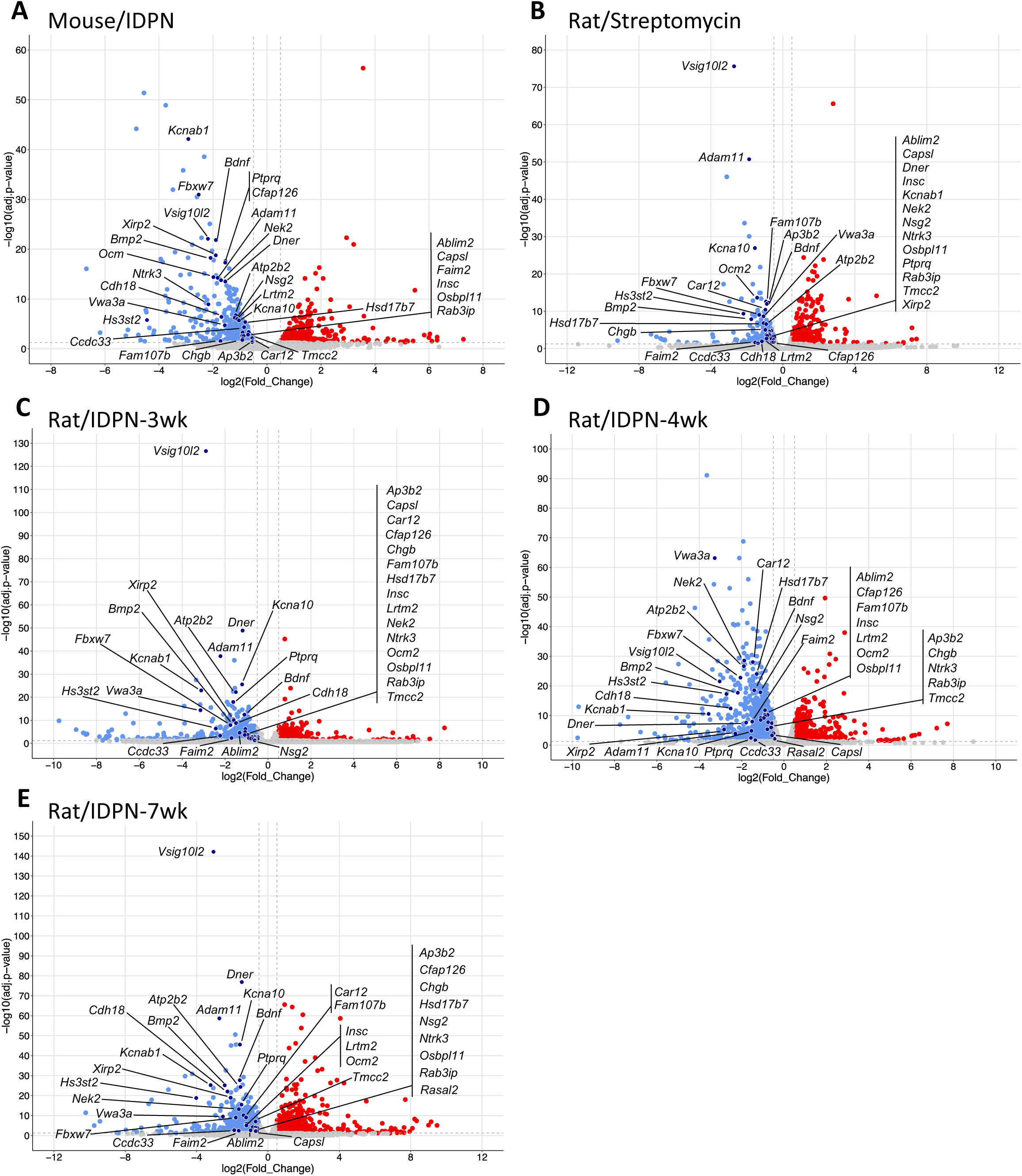
Volcano plots showing the differentially expressed genes (DEG) in control vs. treated animals according to their log(Padj) and log2-Fold_Change values. Data are shown for IDPN in mice **(A)**, streptomycin in rats **(B)**, and IDPN in rats for 3 weeks **(C)**, 4 weeks **(D)** and 7 weeks **(E)**. The control animals were the same in the rat/IDPN-3wk and rat/IDPN-7wk comparisons. The labels indicate the 34 genes found to be downregulated in all five control vs. treated comparisons. N=3/group.

**Table 1.**
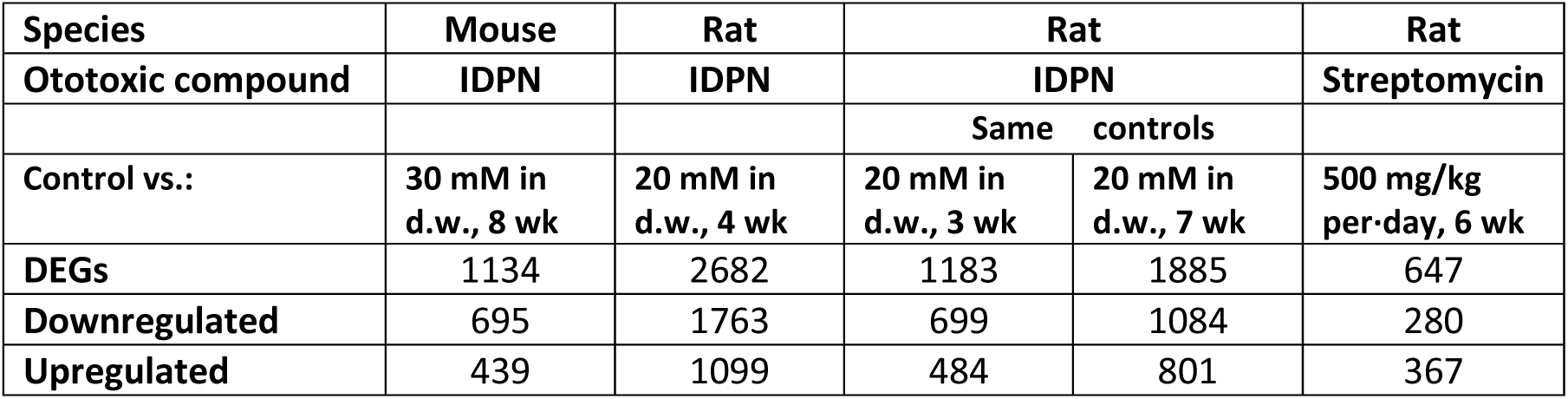
Number of Differentially Expressed Genes between control and treated animals (n=3/group)

### 2.3. RNA-seq: DOWNREGULATION RESPONSES

#### Gene downregulation responses common to all subchronic ototoxicity models

GO analyses of downregulated genes identified from 59 (in rat/IDPN-3wk) to 654 (in rat/IDPN-4wk) Biological Process (BP) terms, from 22 (in rat/STREP) to 157 (in rat/IDPN-4wk) Cell Component (CC) terms, and from 26 (in rat/IDPN-3wk) to 95 (in rat/IDPN-4wk) Molecular Function (MF) terms. To obtain a focused view on the effects of the treatments, we excluded from these lists terms with 1000 or more genes, making them too broad. The resulting lists of enriched terms are provided with their corresponding p-values in Suppl. Tables S1. Comparison of these lists across experimental conditions revealed the presence of terms common in all comparisons. Fig. 4 shows the terms enriched in the five lists of downregulated genes, displayed according to the significance adjusted p-values in the control vs. streptomycin comparison. The most prominent group of BP terms identified by the analysis (Fig. 4A) was that of terms related to cell structure, development, or function of HCs. Thus, highlighted terms in all five BP lists included, among others, “sensory perception of sound” (GO:0007605) and “sensory perception of mechanical stimulus” (GO:0050954), “mechanoreceptor differentiation” (GO:0042490) and “inner ear development” (GO:0048839). In addition, we identified two other conspicuous groups of BP terms associated with the genes downregulated in all exposure conditions. One was that of terms related to ion channels such as “monoatomic ion transmembrane transport” (GO:0034220). This was paralleled by a group of MF terms also identified in all five control vs. treated comparisons (Fig. 4B) that included “monoatomic cation channel activity” (GO:0005261)“ and other related terms. Also enriched were several BP terms related to the synapse such as “chemical synaptic transmission” (GO:0007268), “anterograde trans-synaptic signaling”(GO:098916), “trans-synaptic signaling” (GO:0099537), and “synaptic signaling” (GO:0099536). Another prominent group of GO terms enriched among downregulated genes was that of terms related to the stereocilium. As shown in Fig. 4C, the CC terms identified in all five comparisons included among others “stereocilium bundle” (GO:0032421), “stereocilium” (GO:0032420), “cluster of actin-based cell projections” (GO:0098862), and “actin-based cell projection” (GO:0098858)”.

**Figure 4.**
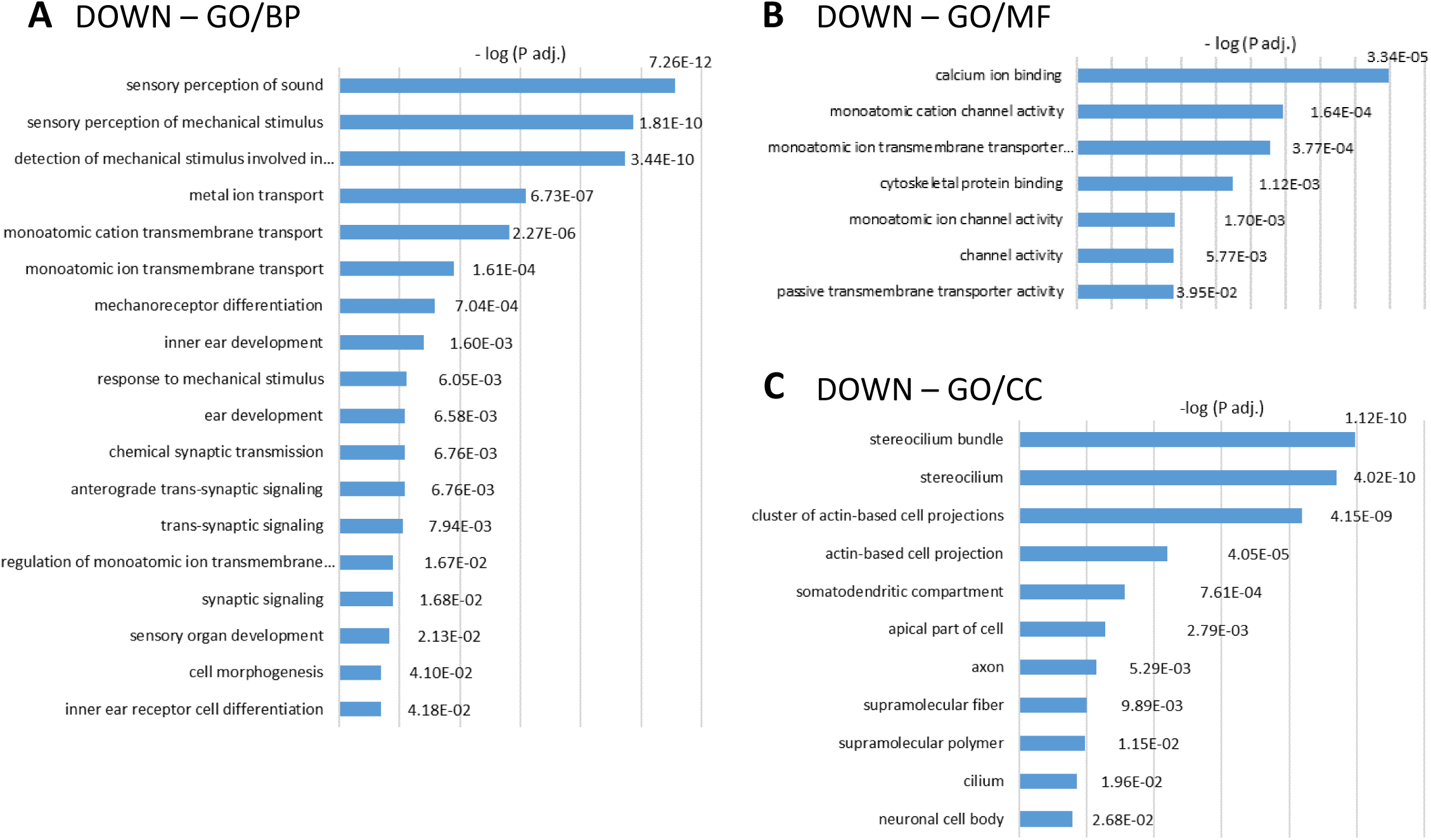
Gene ontology (GO) analyses of the lists of downregulated genes. Graphs show the lists of GO terms significantly enriched in all five control *vs*. treated comparisons: mouse/IDPN, rat/STREP, rat/IDPN-4wk, rat/IDPN-3wk, and rat/IDPN-7wk. Broad terms (> 1000 genes) were excluded. The annotated bars show the –log(Padj) values attained by the corresponding term in the comparison of control *vs*. streptomycin rats. **(A):** Biological process (BP). **(B):** Molecular function (MF). **(C):** Cell component (CC).

A parallel analysis of common gene downregulation response shared by all experimental conditions was performed by crossreferencinging the lists of gene symbols to identify the genes present in all five lists. As shown in the Venn diagram (Fig. 5A), 34 genes were identified as genes downregulated in all experimental models of ototoxicity studied. Among these genes, listed in table 2, there are trophic factors and neurotrophin receptors (*Bdnf*, *Bmp2*, *Ntrk3*), as well as proteins with known roles in terminal differentiation of HCs and stereocilium morphology (*Xirp2*, *Dner*, *Ptprq*, *Cfap126*), cell adhesion (*Cdh18*, *Adam11*), vesicular secretion (*Chgb*), stereocilium calcium pumping (*Atp2b2*) and potassium permeation (*Kcnab1*, *Kcna10*). Fig. 5B shows the expression heat maps of these 34 genes in the 5 control vs. treated comparisons. GO analysis of this list of 34 DEGs identified “mechanoreceptor differentiation” (GO:0042490) and “inner ear development” (GO:0048839) as the only two significantly enriched BP terms (Padj = 4.41E-03 and Padj = 6.98E-03, respectively). As shown in table 2 and discussed below, this results from the fact that 29 of these 34 genes have been previously identified to be selectively expressed by HCs within the rodent or human vestibular sensory epithelium. Thus, the data revealed the downregulation of genes that play an important role in mature HC function and terminal differentiation.

**Figure 5.**
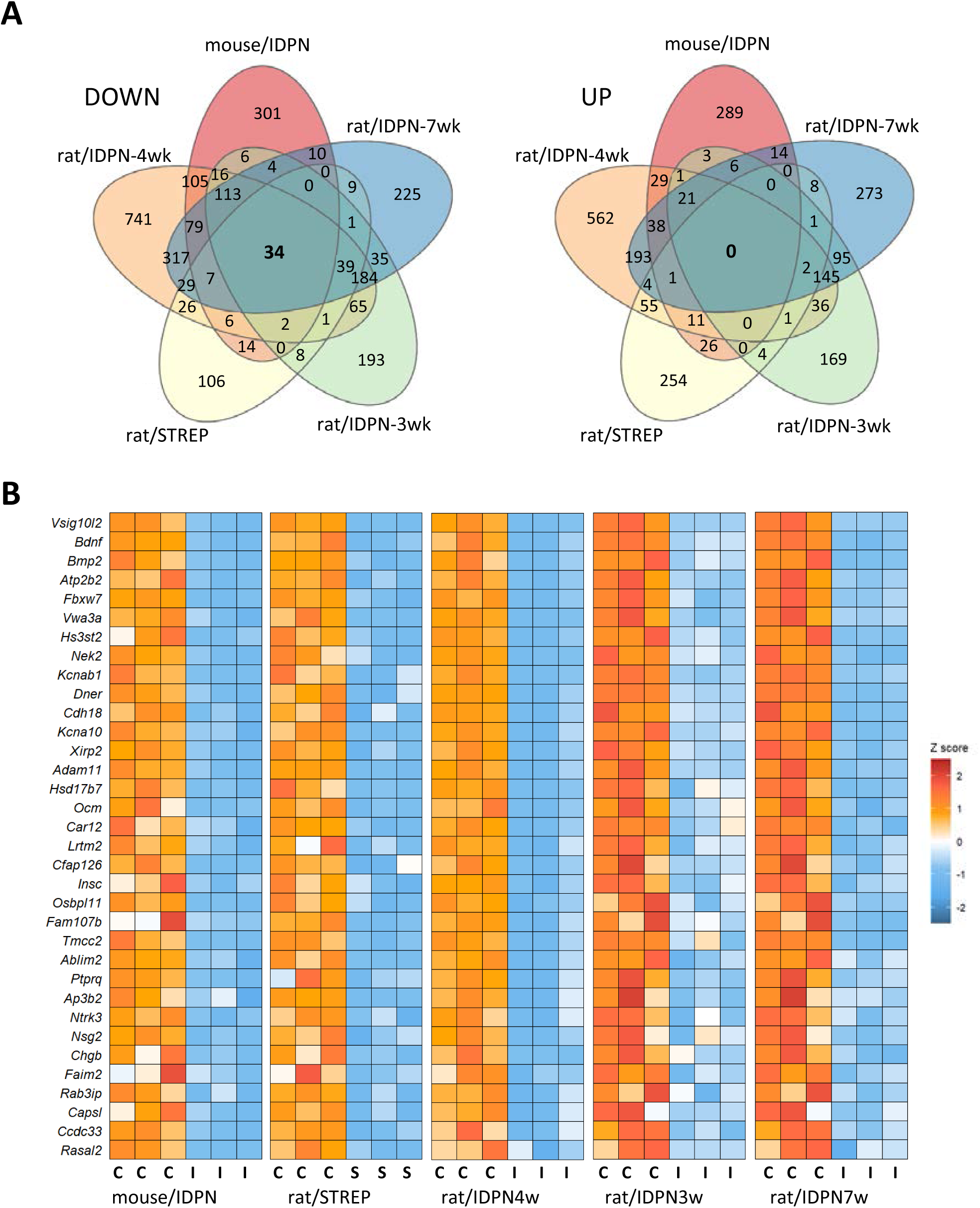
(A) Venn diagrams of downregulated and upregulated genes in the five control vs. treated comparisons in the study: mouse/IDPN, rat/STREP, rat/IDPN-4wk, rat/IDPN-3wk, and rat/IDPN-7wk. Comparison of DEGs lists from each control vs treated comparison identified 34 genes downregulated in rats exposed to streptomycin that were also downregulated in mice and rats exposed to IDPN at any stage of exposure. In contrast, no common gene was identified to be upregulated in all five comparisons. **(B)** Heat maps of the expression of the 34 genes downregulated in all comparisons.

**Table 2.** Genes down-regulated in all models.

| Gene symbol | Name | HC vs SC Selectivity |
| --- | --- | --- |
| **Ablim2** | Actin binding LIM protein family, member 2 | Yes |
| **Adam11** | ADAM metallopeptidase domain 11 | Yes |
| **Ap3b2** | Adaptor related protein complex 3 subunit beta 2 | Yes |
| **Atp2b2** | ATPase plasma membrane Ca<sup>2+</sup> transporting | Yes |
| **Bdnf** | Brain-derived neurotrophic factor | Yes |
| **Bmp2** | Bone morphogenetic protein 2 | Yes |
| **Capsl** | Calcyphosine-like | Yes |
| **Car12** | Carbonic anhydrase 12 | Yes |
| **Ccdc33** | Coiled-coil domain containing 33 | Yes |
| **Cdh18** | Cadherin 18 | Yes |
| **Cfap126** | Cilia and flagella associated protein 126 | - |
| **Chgb** | Chromogranin B | Yes |
| **Dner** | Delta/notch-like EGF repeat containing | Yes |
| **Faim2** | Protein lifeguard 2-like | Yes |
| **Fam107b** | Family with sequence similarity 107, member B | - |
| **Fbxw7** | F-box and WD repeat domain containing 7 | Yes |
| **Hs3st2** | Heparan sulfate-glucosamine 3-sulfotransferase 2 | Yes |
| **Hsd17b7** | Hydroxysteroid (17-beta) dehydrogenase 7 | Yes |
| **Insc** | INSC, spindle orientation adaptor protein | Yes |
| **Kcna10** | Potassium voltage-gated channel subfamily A member 10 | Yes |
| **Kcnab1** | Potassium voltage-gated channel subfamily A member regulatory beta subunit 1 | Yes |
| **Lrtm2** | Leucine-rich repeats and transmembrane domains 2 | Yes |
| **Nek2** | NIMA-related kinase 2 | Yes |
| **Nsg2** | Neuronal vesicle trafficking associated 2 | Yes |
| **Ntrk3** | Neurotrophic receptor tyrosine kinase 3 | - |
| **Ocm/Ocm2** | Oncomodulin | Yes |
| **Osbpl11** | Oxysterol binding protein-like 11 | - |
| **Ptprq** | Protein tyrosine phosphatase, receptor type, Q | Yes |
| **Rab3ip** | RAB3A interacting protein | Yes |
| **Rasal2** | RAS protein activator like 2 | - |
| **Tmcc2** | Transmembrane and coiled-coil domain family 2 | Yes |
| **Vsig10l2** | V-Set and immunoglobulin domain containing 10 like 2 | Yes |
| **Vwa3a** | von Willebrand factor A domain containing 3A | Yes |
| **Xirp2** | Xin actin-binding repeat containing 2 | Yes |

Yes: Gene selectively expressed in HC versus supporting cells, according to literature data (Wu et al., 2013; Burns et al., 2015; Scheffer et al., 2015; Sadler et al., 2020; Wang et al., 2024).

For additional identification of genes downregulated both after IDPN or streptomycin exposure, we compared the gene lists of the rat/STREP and rat/IDPN-4 wk experiments. The 4-week time point of exposure to IDPN was selected for this additional comparison with the streptomycin data because it likely corresponded to a more similar level of toxicity than that of IDPN rats examined after shorter (3 weeks) or longer (7 weeks) times of exposure. In addition to the 34 genes listed in Table 2, 108 additional genes were identified, resulting in a total of 142 genes similarly downregulated in the vestibular epithelium of streptomycin and IDPN rats (Fig. 6A). In this extended list, several additional genes that encode important HC proteins were recognized. Thus, the list contains genes for the motor protein *Myo7a*, the calcium binding protein *Pvalb*, the tyrosine phosphatase *Ptprz1* and a component of the mechanotransduction complex *Lhfpl5*, all of which are known to play a prominent role in the physiology of HCs. Consequently, the GO analysis of this list identified “sensory perception of sound” (GO:0007605), with Padj = 5.56E-07, and “stereocilium bundle” (GO:0032421), with Padj = 2.34E-05, as the BP and CC terms most significantly enriched, respectively (Fig. 6B). An enrichment value of 1.0 in the MF term “calcium-dependent ATPase activity” (GO:0030899) revealed the decreased expression of both the Atp2b2 and Atp2b3 genes, which encode the PMCA and SERCA2 proteins.

**Figure 6.**
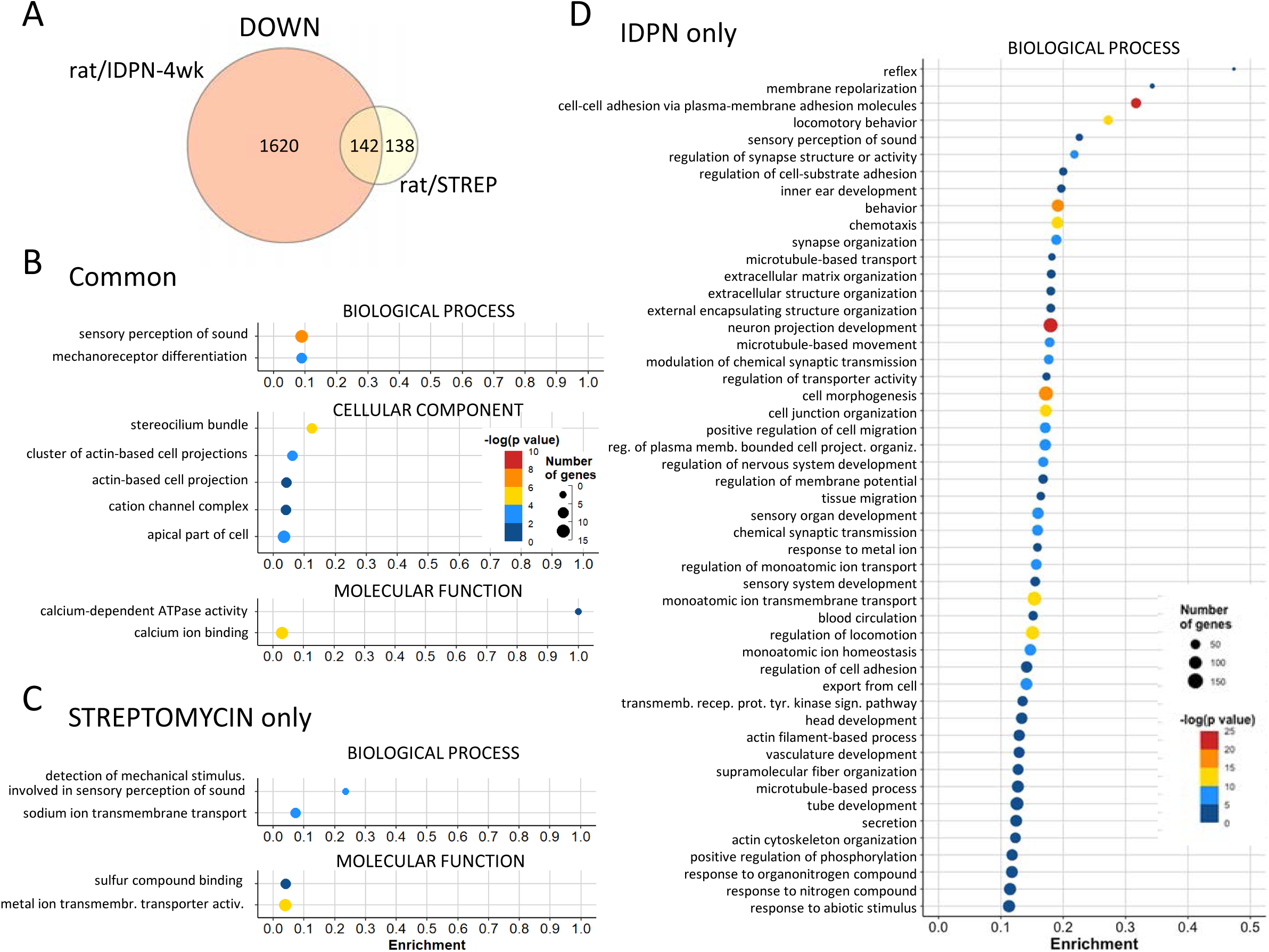
Common and unique downregulation responses in the rat/STREP and rat/IDPN-4wk groups. **(A)** Venn diagram showing the numbers of downregulated genes in one, the other, or both experiments. **(B, C, D)** Results of GO analysis of the three lists of genes downregulated after either treatment (**B**), after streptomycin only (**C**), or after IDPN only (**D**). In all three cases, the original list was reduced by REVIGO analysis of redundancy and elimination of broad terms (> 1,000 genes). All terms identified after reduction are shown in **(B)** and **(C)**; only BP terms are shown in **(D)**.

For an alternative approach to identify common downregulation responses elicited by subchronic ototoxicity exposure, we performed the threshold-free GSEAS to identify gene sets enriched in control animals. In each experimental condition, only one or two gene sets were enriched in control compared to treated samples at FDR < 0.25, and none of them were common across models (data not shown). Therefore, no hallmark gene set was downregulated in all subchronic ototoxicity models.

Together, these comparisons lead to the conclusion that enduring ototoxic stress induces a decrease in the expression of HC-specific genes. An apparent decrease in expression could result from a reduced number of HCs present in the sequenced samples. However, based on previous assessment of these exposure models under study (Sedó-Cabezón et al., 2015; Greguske et al., 2019; Maroto et al., 2023), the data in Fig. 1 E-G indicate that only a limited loss of HCs affected most of the sequenced samples. So the observed large decreases in RNA counts resulted from the downregulation of gene expression, as outlined above. This conclusion was corroborated by the validation data shown below. The observed downregulation response is shared by exposure to IDPN and streptomycin, occurs in both rats and mice, and begins in the early stages of chronic ototoxicity in association with the initial loss of vestibular function. We also concluded (see below) that this response widens as ototoxic stress progresses at longer times of exposure.

#### Compound-specific gene downregulation responses in vestibular epithelia after subchronic ototoxicity

In addition to identifying genes that respond similarly in all exposure models, we used the data to identify genes that responded differently with one or the other toxic compound. Using the rat/STREP and rat/IDPN-4wk datasets, we identified the genes downregulated in streptomycin rats but not in IDPN rats, and those downregulated by IDPN but not streptomycin (Fig. 6A). The 138 rat/STREP-only genes did not reveal any prominent differential response; they were mostly genes that were not significantly downregulated in the rat/IDPN-4wk samples but they were related to GO terms that were also downregulated by IDPN exposure, such as “sodium ion transmembrane transport” (GO:0035725) (Fig. 6C). The analysis of the 1620 genes unique to the rat/IDPN-4wk downregulation response identified many GO terms also identified in the rat/STREP experiment, such as “sensory perception of sound” (GO: 0007605), as explained above. However, other terms were apparently exclusive to the IDPN effect. A particularly consistent difference was that of terms related to cell adhesion, such as “cell-cell adhesion via plasma-membrane adhesion molecules” (GO:0098742), as many cadherins, protocadherins, CAMs and other adhesion factors were downregulated in the rat/IDPN-4wk but not in the rat/STREP samples (Fig. 6D). The effect of IDPN toxicity on cell adhesion was also present in the other IDPN experiments, and robust significance of many related terms, including “cell-cell adhesion via plasma-membrane adhesion molecules”, was present in mouse/IDPN, rat/IDPN-3wk and rat/IDPN-7wk versus control comparisons.

#### Downregulation responses in the late stages of subchronic ototoxicity

In the advanced stages of subchronic IDPN toxicity, HCs alter their morphology and start to extrude from the sensory epithelium into the endolymphatic cavity. Abnormal and extruding HCs accumulate in the epithelium, indicating that the process is characterized by considerable synchrony and slow progression (Seoane et al., 2001a,b; Sedó-Cabezón et al., 2015). In contrast, our previous data (Maroto et al., 2023) suggest that this form of HC death may operate in streptomycin rats, but do so in a much more non-synchronic and fast-peaced manner, meaning partly extruded cells do not accumulate in the epithelium. To evaluate whether late staged damage (i.e. extruding or pre-extruding) to sensory HCs present a downregulated response different from that at an earlier stage, we focused on the genes downregulated in rat/IDPN-7wk specimens but not in rat/IDPN-3wk or rat/STREP specimens. The list included 631 genes (Fig. 7A) that, according to GO analysis, were associated with 95 BP terms. These terms showed a significant overlap with terms associated with genes that were similarly downregulated in all 5 models. For example, several terms related to HCs, grouped by REVIGO under the term “inner ear development” (GO:0048839), were significantly altered (Fig. 7B). Therefore, more HC-specific genes were downregulated at later time points in IDPN ototoxicity than at earlier stages. Among them, there were genes such as *Atoh1* and *Pou4f3*, key transcription factors in HC differentiation, or *Cdh23* and *Ush1c*, which encode the tip-link proteins cadherin-23 and harmonin, respectively. Similarly, the cluster of enriched terms under the term “synapse organization” (GO:0050808) matched the terms enriched in all five comparisons.

**Figure 7.**
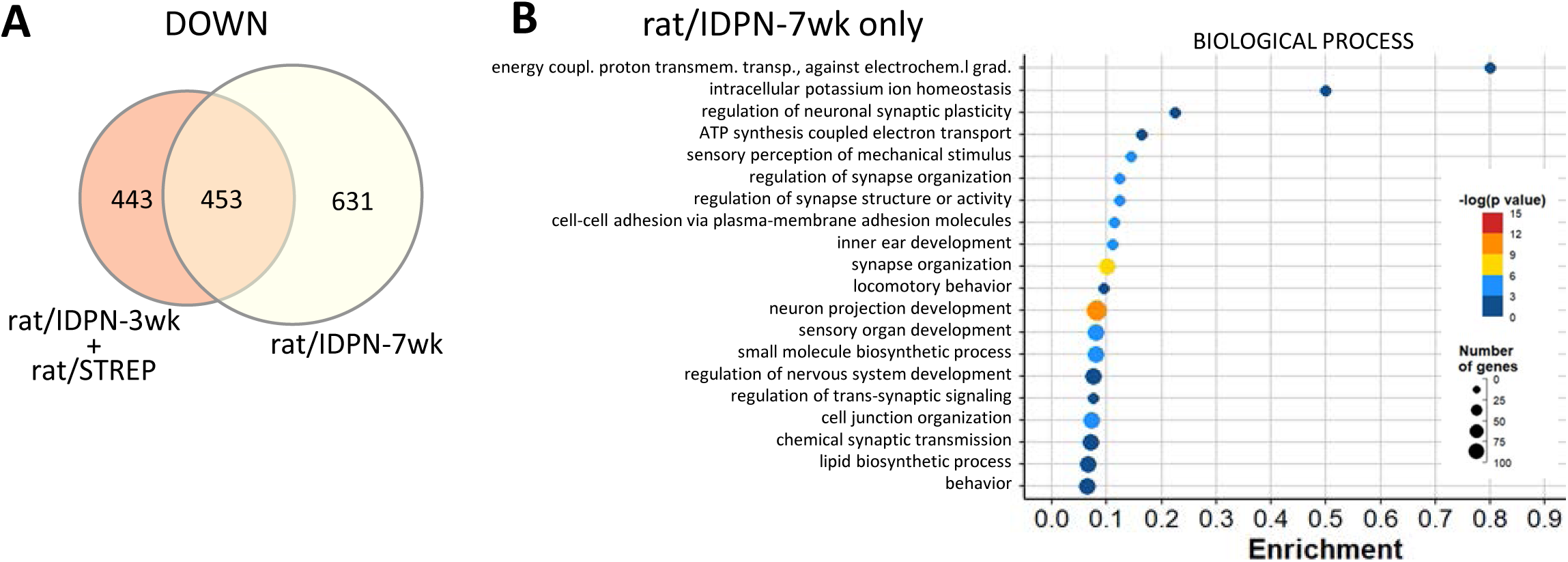
Comparison of downregulation responses in the most affected (rat/IDPN-7wk) *vs.* the least affected (rat/IDPN-3wk + rat/STREP groups). **(A)** Venn diagram showing the numbers of downregulated genes in one, the other, or both groups of experimental conditions. **(B)** Results of the GO analysis of the 631 genes downregulated in IDPN-7wk rats but not in IDPN/3-wk or STREP rats. Only the BP terms are shown, after reduction of the original list by REVIGO analysis and elimination of broad terms (> 1,000 genes).

Therefore, these observations further support the conclusion stated above that the downregulation response shared by all ototoxicity models widens after longer times of exposure. In addition, other terms derived from the list of genes downregulated after 7 but not 3 weeks of exposure to IDPN overlapped with those shared by all IDPN models, such as terms related to cell adhesion, indicating that additional genes of the same domain significantly reduced expression in later stages of toxicity. Nevertheless, at least one cluster of BP terms enriched in the list of genes downregulated in rat/IDPN-7wk samples but not in rat/IDPN-3wk or rat/STREP samples represented a unique addition of late IDPN toxicity. REVIGO selected the term “ATP synthesis coupled electron transport” (GO:0042773) to represent this cluster. The downregulated genes associated with this term were nuclear (e.g.: *Ndufs4*) and mitochondrially-encoded (e.g.: *mt-Nd4*). So, a qualitative difference between late (rat/IDPN-7wk) and early (rat/IDPN-3wk or rat/STREP) ototoxicity appears to be downregulation of genes involved in mitochondrial respiration.

### 2.4. RNA-seq: UPREGULATION RESPONSES

#### Common gene upregulation responses in all subchronic ototoxicity exposure models

GO analyses of upregulated genes identified from 61 (in rat/IDPN-3wk) to 457 (in rat/IDPN-7wk) BP terms, from 11 (in rat/STREP) to 59 (in rat/IDPN-7wk) CC terms, and from 4 (in rat/IDPN-3wk) to 51 (in rat/IDPN-4wk) MF terms that were significantly enriched (Suppl. Tables S1). Comparison of these lists of terms across the 5 experimental evaluations identified only four broad CC or MF terms in common (GO:0005737, “cytoplasm”; GO:0005829, “cytosol”; GO:0003824, “catalytic activity”; GO:0005515, “protein binding”). Among BPs, 23 terms were significantly enriched after all ototoxic treatments. This list of terms, shown in Fig. 8 according to the p-adj values in the streptomycin experiment, was only moderately informative, as all of these terms were broad, including more than 1000 genes. Nevertheless, several of them were related to stress responses and responses to chemicals. Comparison of the original lists of upregulated genes did not identify any gene that was more expressed than in controls in all 5 comparisons (Fig 5A). However, if the mildest toxicity condition (rat/IDPN-3wk) was excluded, a single gene was found to be upregulated in the other 4 conditions and this gene was *Atf3*. For a broader view of the upregulation responses shared by IDPN and streptomycin, we compared the DEG lists from the rat/IDPN-4wk and rat/STREP experiments and obtained a list of 74 common upregulated genes (Fig 9A). GO analysis of this list (Fig. 9B) identified “positive regulation of transcription from RNA polymerase II promoter in response to stress” (GO:0036003) as the most significantly associated BP term, because of the presence of *Atf3*, *Atf4*, *Cebpb* and *Notch1* in the list. The list also contained five additional genes that encode components of the RNA polymerase II transcription regulatory complex (*Esf4*, *E2f6*, *Vdr*, *Taf5l*, *LOC103689931*). These results suggest that chronic ototoxic stress caused by various types of toxic compounds (IDPN and streptomycin) can trigger, in the vestibular epithelium of both rats and mice, a common stress response involving ATF3, ATF4, and Cebpb. However, this transcriptional response seems to start later than the downregulation of HC-marker genes identified above during progressive toxicity.

**Figure 8.**
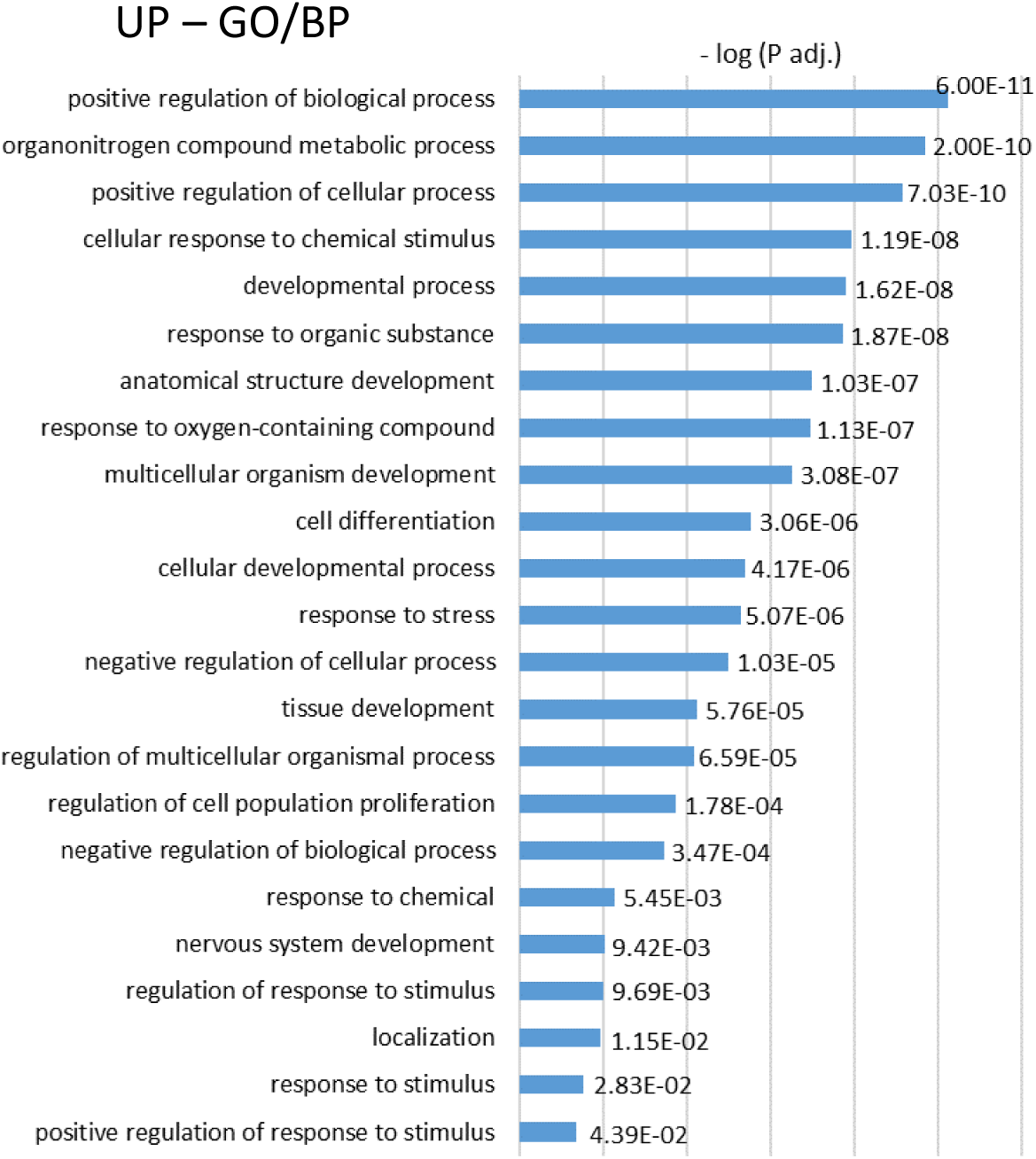
Gene ontology (GO) analyses of the lists of upregulated genes. The graph shows the list of BP terms significantly enriched in all five control *vs*. treated comparisons: mouse/IDPN, rat/STREP, rat/IDPN-4wk, rat/IDPN-3wk and rat/IDPN-7wk. All terms were broad (> 1000 genes). The annotated bars show the –log(Padj) values attained by the corresponding term in the comparison of control *vs.* streptomycin rats.

**Figure 9.**
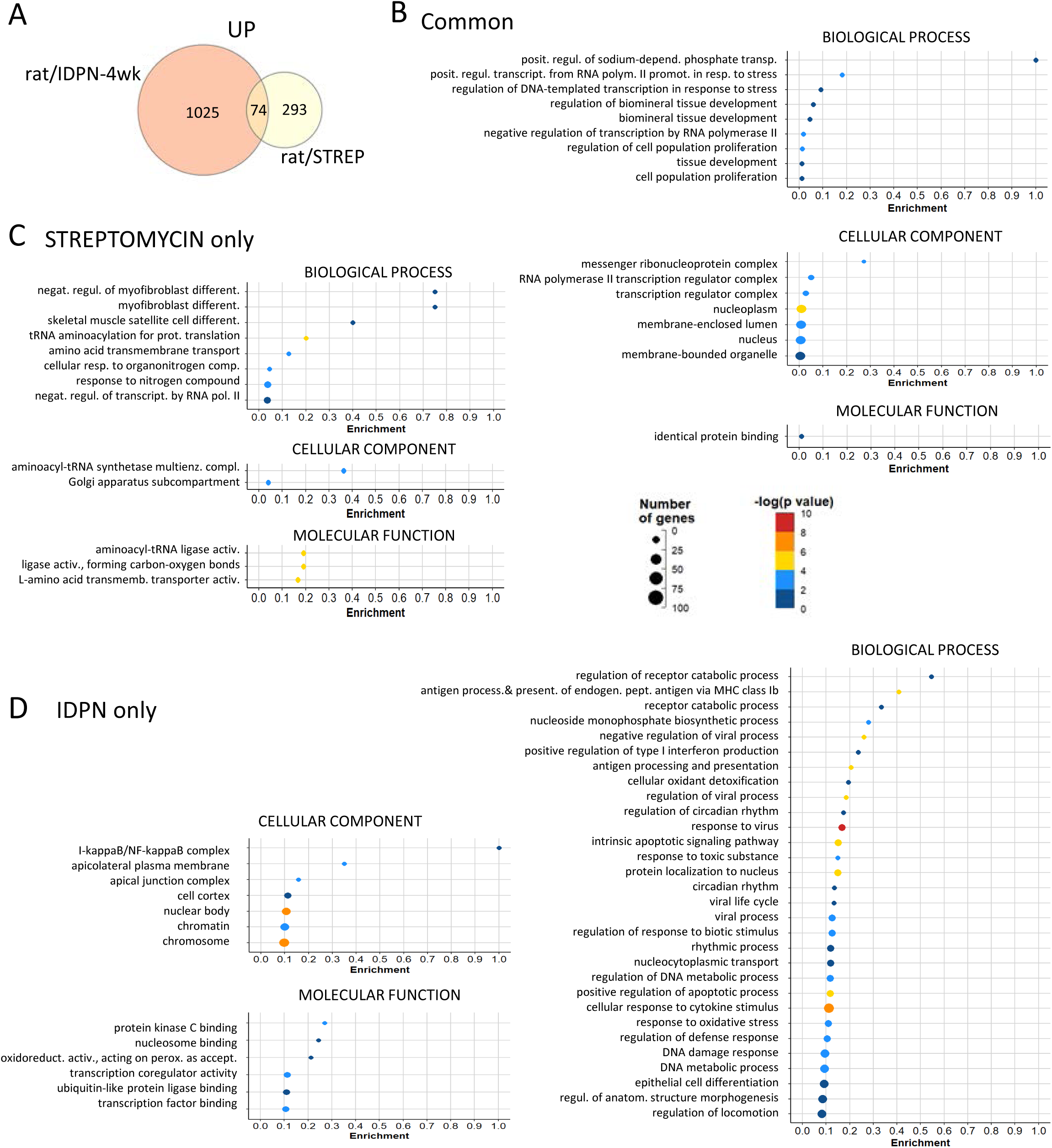
Common and unique upregulation responses in the rat/STREPTO and rat/IDPN-4wk groups. **(A)** Venn diagram showing the numbers of genes upregulated in one, the other, or both experiments. **(B, C, D)** Results of GO analysis of the three lists of genes upregulated after either treatment (**B**), only after streptomycin (**C**), or only after IDPN (**D**). In all three cases, the original list was reduced by REVIGO analysis of redundancy and elimination of broad terms (> 1,000 genes).

By GSEAS, there was a clear difference between IDPN-upregulated gene sets and streptomycin-upregulated gene sets, with no overlap at FDR < 0.05 (Table 3). However, streptomycin samples showed marginal enrichment in the gene sets of the p53 pathway (FDR=0.055) and MYC_Targets_V1 (FDR=0.060), and these were significantly enriched in the IDPN groups except for the IDPN/3wk group.

**Table 3.** Hallmark gene sets identified by GSEA as significantly enriched (FDP q-value < 0.05) in treated vs control samples in the five comparisons in the study.

| Comparison | Mouse/IDPN | Rat/IDPN-4wk | Rat/IDPN-3wk | Rat/IDPN-7wk | Rat/STREP |
| --- | --- | --- | --- | --- | --- |
|  |  |  | (same controls) |  |  |
| Control vs.: | 30 mM in d.w., 8 wk | 20 mM in d.w., 4 wk | 20 mM in d.w., 3 wk | 20 mM in d.w., 7 wk | 500 mg/kg per-day, 6 wk |
| 1st significant gene set (FDR q-value) | Interferon-alpha response (0.000) | Interferon-alpha response (0.000) | MYC_Targets_V1 (0.000) | MYC_Targets_V1 (0.000) | Unfolded protein response (0.000) |
| 2 | Interferon-gamma response (0.000) | Interferon-gamma response (0.000) | Interferon-alpha response (0.000) | Interferon-alpha response (0.004) | WNT_beta_catenin signaling (0.048) |
| 3 | TNFa signalling via NFkB (0.000) | IL6_JAK_Stat3_signaling (0.001) | IL6_JAK_Stat3 signaling (0.010) | IL6_JAK_Stat3 signaling (0.012) | *TNFa signalling via NFkB (0.051)* |
| 4 | Inflammatory response (0.002) | E2F Targets (0.001) | Apical surface (0.020) | E2F Targets (0.014) | *p53 Pathway (0.055)* |
| 5 | IL6_JAK_Stat3 signaling (0.003) | MYC_Targets_V1 (0.000) | E2F Targets (0.016) | UV_response_up (0.015) | *MYC_Targets_V1 (0.060)* |
| 6 | Allograft rejection (0.007) | MYC_Targets_V2 (0.000) | UV_response_up (0.019) | Apical surface (0.016) |  |
| 7 | TGF-beta signaling (0.010) | TNFa signalling via NFkB (0.001) | Interferon-gamma response (0.026) | Interferon-gamma response (0.023) |  |
| 8 | PI3K_Akt_mTOR signaling (0.015) | G2M Checkpoint (0.002) | Apoptosis (0.025) | Apoptosis (0.026) |  |
| 9 | G2M Checkpoint (0.049) | Apoptosis (0.004) | DNA_Repair (0.027) | p53 Pathway (0.024) |  |
| 10 |  | p53 Pathway (0.008) | MYC_Targets_V2 (0.025) | Allograft rejection (0.023) |  |
| 11 |  | UV_response_up (0.009) | p53 Pathway (0.023) | MYC_Targets_V2 (0.021) |  |
| 12 |  | Allograft rejection (0.024) | Allograft rejection (0.025) | DNA_Repair (0.025) |  |
| 13 |  | Hypoxia (0.036) | Xenobiotic metabolism (0.026) | Xenobiotic metabolism (0.025) |  |
| 14 |  |  | Hypoxia 0.032 | Reactive_oxygen_species pathway (0.032) |  |
| 15 |  |  | Reactive_oxygen_species pathway (0.033) | Hypoxia (0.032) |  |
| 16 |  |  | WNT_beta_catenin signaling (0.043) | Coagulation (0.043) |  |
| 17 |  |  | Coagulation (0.043) |  |  |

#### Compound-specific gene upregulation responses in vestibular epithelia after subchronic ototoxicity

As previously done for downregulation responses, we compared the list of rat/STREP and rat/IDPN-4wk genes to identify upregulation responses that may be unique to streptomycin and IDPN toxicities. Thus, we identified a list of 293 genes that were upregulated after streptomycin but not after 4 weeks of IDPN in rats (Fig. 9A). GO analysis of this list followed by REVIGO reduction and exclusion of broad terms (term_size > 1000) resulted in a list of 8 BP, 2 CC and 3 MF terms (Fig. 9C). This list included “tRNA aminoacylation for protein translation” (GO:0006418) as the most significantly enriched BP term along with the CC term “aminoacyl-tRNA synthetase multienzyme complex” (GO:0017101), and the MF term “aminoacyl-tRNA ligase activity” (GO:0004812). We noticed significant upregulation of 9 genes that encode tRNA aminoacylation proteins such as *Yars1* and *Lars1*. Other significant terms were “amino acid transmembrane transport” (BP, GO:0003333) and similar terms, due to the presence of several amino acid transporters of the Slc family, such as the high affinity cationic amino acid transporter, *Slc7a1*, and the glutamate/neutral amino acid transporter, *Slc1a4*. Therefore, subchronic streptomycin, but not IDPN, induced the expression of genes involved in protein synthesis. By GSEA, streptomycin samples, but none of the IDPN groups, showed a highly significant enrichment of the “unfolded protein response” hallmark gene set (Table 3).

Up to 1025 genes were upregulated in IDPN-4wk rats but not after streptomycin (Fig. 9A). After GO analysis, REVIGO downsizing and exclusion of broad terms, many BP terms related to stress, immune, defence, and anti-viral responses were identified (Fig 9D). This resulted from the presence of interferon-responsive genes and immune-related genes in the lists, such as *Stat1*, *Stat2*, *Stat3*, *Irf7*, *Irf9*, *Oas2*, *Igtp*, *Isg15* and many others. When we evaluated upregulation in all control *vs* treated comparisons, we observed that this immune-related response was also present in all IDPN groups (mouse/IDPN, rat/IDPN-3wk, rat/IDPN-4wk, and rat/IDPN-7wk). In good agreement with these GO analyses, GSEA identified the enrichment of several related hallmark sets of genes in common in all four IDPN *vs* control comparisons but not in the streptomycin *vs* control comparison (Table 3). Specifically, the gene sets “interferon-alpha response”, “interferon-gamma response”, “IL6-JAK-STAT3 signalling” and “allograft rejection” were significantly enriched (FDR <0.05) after IDPN but not after streptomycin exposure.

#### Upregulation responses in late stages of subchronic ototoxicity

As explained above for the downregulation responses, we searched for late stage ototoxicity upregulation responses by studying the list of genes upregulated in rat/IDPN-7wk specimens but not in rat/IDPN-3wk or rat/STREP models. This list contained 518 genes (=245+273, upper right in Fig.10A). The result of the GO analysis followed by the REVIGO reduction highlighted the upregulation of anti-viral response genes, revealing that the lists of upregulated interferon-stimulated genes were longer in late stage than in early stage IDPN exposure conditions. Thus, genes such as *Stat1*, *Stat2*, *Irf7*, *Irf9*, *Igtp*, and *Isg15* were upregulated in the rat/IDPN-7wk but not in the rat/IDPN-3wk samples. Nevertheless, other genes related to anti-viral responses and interferon signalling, such as *Usp18*, *Oas1a*, *Anxa1*, *Ifitm1*, *Ifi27*, *Ifitm3*, and *Eif2ak2,* were already upregulated in the sensory epithelia of IDPN-3wk rats. For an illustration of the core response to advanced IDPN ototoxicity, we selected the 245 genes that were upregulated in both the rat/IDPN-7wk samples and the rat/IDPN-4wk or mouse/IDPN samples, but not in the rat/IDPN-3wk or rat/STREP specimens. Cluster analysis of protein-protein interactions of this list by STRING (von Mering et al., 2003; Szklarczyk et al., 2023), using an MLC inflation parameter of 2, identified a cluster of 53 proteins shown in Fig. 10B as the main result. The top BP descriptor of this cluster was “Defense response to virus” (GO:0051607; FDR = 3.10E-30) and many related terms associated with it. These included, for example, “Response to virus” (GO:0009615; FDR =1.38E-33), “Negative regulation of viral process” (GO:0048525; FDR = 6.28E-18), and “Response to interferon-beta” (GO:0035456; FDR = 1.12E-13), indicating that progression of IDPN ototoxicity consistently causes a broadening anti-viral-like expression response.

**Figure 10.**
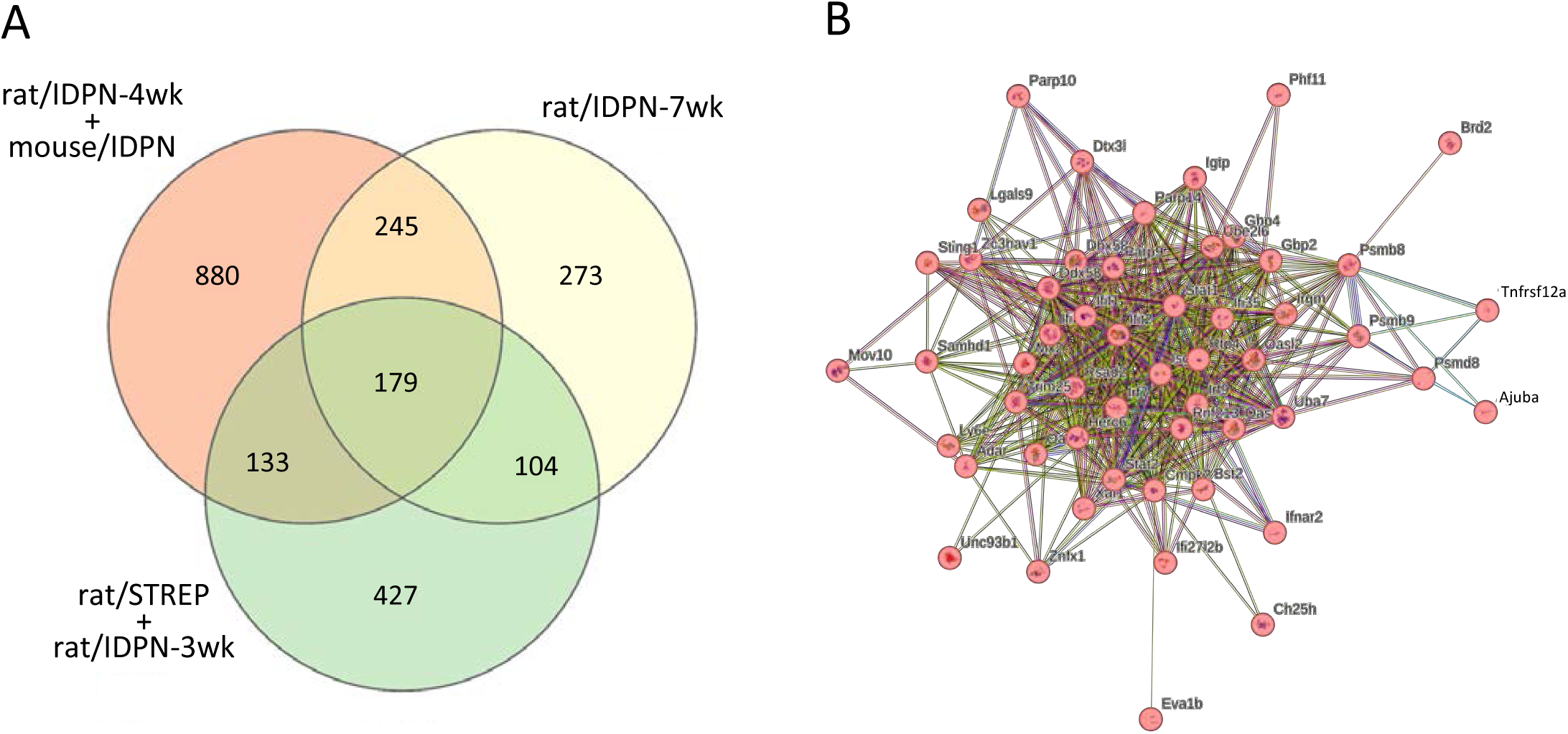
Comparison of upregulation responses in the most affected (rat/IDPN-7wk) vs. the least affected (rat/IDPN-3wk + rat/STREP) vs. the advanced toxicity (rat/IDPN-4k + mouse/IDPN) groups. **(A)** Venn diagram showing the numbers of genes upregulated in each group of experimental conditions and their intersections. **(B)** Cluster of 53 functionally associated proteins identified by STRING analysis of the list of 245 core genes upregulated in advanced IDPN ototoxicity, defined as genes upregulated in both the rat/IDPN-7wk samples and the rat/IDPN-4wk or mouse/IDPN samples, but not in the rat/IDPN-3wk or rat/STREP specimens.

### 2.5. VALIDATION STUDIES

#### The gene downregulation response triggered by subchronic ototoxicity is associated with decreased expression of the corresponding proteins in HCs

As explained above, all ototoxic treatments caused a reduction in the number of transcripts from 34 genes, 29 of which have been demonstrated to be selectively expressed in HCs in human or rodent vestibular epithelium. The epithelia used for sequencing were affected by no or only limited loss of HC (sections 2.1. and 2.3.), so we can conclude that vestibular HCs under sustained toxic stress downregulate the expression of genes encoding proteins that characterize their terminal differentiation and are key to their transducing roles. To further support this conclusion, we used immunofluorescent labelling and confocal microscopy to compare the expression of some of these proteins in samples from control and treated animals in the rat/IDPN-4wk and rat/STREPTO experiments and in vestibular epithelia exposed to streptomycin *in vitro*. *Atp2b2*, which encodes the plasma membrane calcium transporting ATP-ase 2 (PMCA2), was robustly downregulated. This calcium transporter is expressed in the stereocilia of auditory and vestibular HCs and extrudes calcium into the endolymphatic cavity (Fettiplace and Nam, 2019). In the rat vestibule, we have recently described that PMCA2 is expressed by HCIs but not by HCIIs (Borrajo et al., 2024). We found that rats exposed to 20 mM of IDPN for 4 weeks had a 72% reduction (t_5df_ = 7.812, p=0.001) in fluorescence intensity of the PMCA2 label in the stereocilia bundles of the utricle (Fig. 11A, Da). Similarly, streptomycin rats (500 mg/kg/day x 6 weeks) showed an 84% reduction (t_6df_= 11.70, p<0.001) in PMCA2 expression in utricular stereocilia (Fig. 11B, Db).

**Figure 11.**
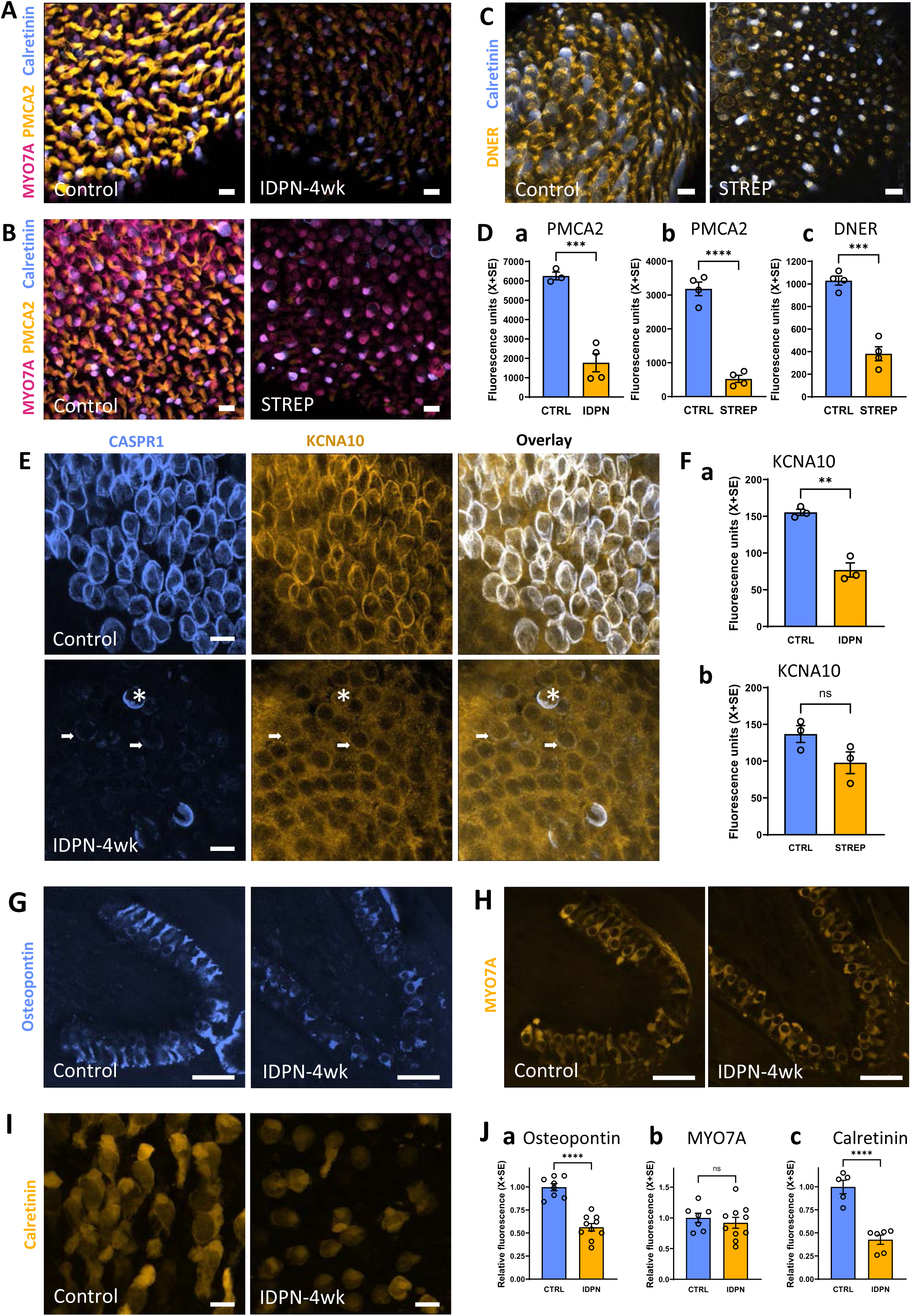
Effect of subchronic ototoxicity on the expression of PMCA2, DNER, KCNA10, osteopontin, calretinin, and MYO7A in vestibular sensory epithelium assessed by immunofluorescent labelling and confocal microscopy. **(A-B)** Images of whole mount utricles from control and treated rats showing the effects of IDPN **(A)** or streptomycin **(B)** on PCMA2 expression in stereocilia bundles. Images are maximum intensity projections of a stack of 6 planes (2 μm total thickness). **(C)** Expression of DNER in the neck region of crista HCs in control and streptomycin rats. Images are maximum intensity projections of a stack of 4 planes (1.2 μm total thickness). **(D)** Quantitative analyses of PCMA2 and DNER expression. Bars show mean +/- SEM fluorescence intensity values. Each point is from an individual animal, averaging values from > 180 hair bundles (**a and b**) or > 400 HCs (c) per animal. *** : p<0.001, ****: p<0.0001, Student’s t-test. (**E**) Immunolabelling of CASPR1 and KCNA10 in a control crista (upper row) and in a crista of a rat exposed to IDPN for 4 weeks (bottom row). In control tissue, both proteins are located in the calyceal junction area. After IDPN, most afferent calyces show very reduced expression of CASPR1 (arrows), while a few retain a large amount of label (asterisk). The corresponding HCs show a marked depletion of the KCNA10 label in the calyceal junction, although the non-specific label shown by supporting cells with this antibody (Martin et al., 2024) persisted. **(F)** Quantitative analysis of the effect of chronic IDPN **(a)** and streptomycin **(b)** on KCNA10 expression in the calyceal junction area. Bars show mean +/- SEM fluorescence intensity values. Each point is from an individual animal, averaging values from > 400 HCs per animal. ** : p<0.01, NS: nonsignificant, Student’s t-test. **(G and H).** Ostepontin and MYO7A labels in sections of cristae from control rats and IDPN-4wk rats. **(I)** Calretinin label in HCII of the periphery of the utricles of control and IDPN-4wk rats. **(J)** Quantitative analyses of the effect of chronic IDPN on the expression of osteopontin, MYO7A, and calretinin. Bars show mean +/- SEM fluorescence intensity values. Each data point is from an individual animal, averaging values of 30 HCs per animal. **** : p<0.0001, ns: nonsignificant, Student’s t-test. Scale bars = 10 μm in A, B, C, E, and H; 25 μm in G and I.

Another downregulated HC-specific gene was the *Dner* gene, which encodes the Delta/Notch-like epidermal growth factor-related receptor (DNER). DNER has been shown to be necessary for proper maturation and orientation of the stereocilia bundle (Kowalik and Hudspeth, 2011). After IDPN exposure, the crista HCs from exposed rats showed a 39 % reduction in DNER expression (t_5df_= 2.96, p=0.031; n=3 control, 4 IDPN) (not shown). A larger effect was recorded after streptomycin exposure, with a 63% reduction in DNER immunoreactivity in crista HCs (t_6dg_=8.73, p<0.001) (Fig. 11C, Dc).

A third protein whose expression was assessed by immunofluorescent confocal microscopy was the potassium voltage-gated channel subfamily A member 10 (KCNA10, also known as Kv1.8), encoded by the *Kcna10* gene. Mutations in this channel cause vestibular impairment (Lee et al., 2013) and recent data have shown its strong expression in the area of the calyceal junction (Martin et al., 2024). In response to subchronic IDPN, the expression of KCNA10 in the calyces was significantly reduced by 50% (t_4df_=7.52, p=0.00167) (Fig. 11E, Fa). After streptomycin treatment, the mean fluorescence value in the calyces of treated animals was 29% lower than the mean in controls (97.9 ± 14.7 vs 136.9 ± 11.5), but this difference did not attain statistical significance (t_4df_=2.091, P=0,104) (Fig 11Fb).

Beyond the list of 34 genes downregulated in all exposed groups, we also aimed to evaluate other HC-specific genes that were downregulated in several exposure conditions and for which reliable antibodies were available. Thus, we assessed anti-osteopontin, anti-calretinin, and anti-MYO7A immunolabelling in rat/IDPN-4wk samples because their encoding genes were down-regulated in this group (*Spp1* [log2-Fold_Change=-1.84; Padj=2.94E-14]; *Calb2* [log2-Fold_Change=-2.17; Padj=9.75E-22]; *Myo7a* [log2-Fold_Change=-0.64; Padj=9.67E-09], respectively). As shown in Fig. 11 G-J, a decrease in immunolabel was recorded for osteopontin (HCI, t_16df_= 7.65, p=0.000) and calretinin (HCII, t_9df_=6.7, p=0.000) but not for MYO7A (all HCs, t_15df_=0.627, p=0.27). We had previously recorded a significant decrease in MYO7A expression in streptomycin-exposed rats compared to control rats (Maroto et al., 2023).

Overall, the immunohistochemical data confirmed that the decreases in the expression of HC-specific genes revealed by the RNA-seq data are associated with decreases in protein expression and are not an artefact resulting from HC loss.

#### *Vsig10l2* is a new HC-specific gene strongly downregulated during HC stress

Among the 34 genes downregulated in all 5 groups of rats exposed to ototoxicity, the *Vsig10l2* gene (previously identified as *Gm1113* in the mouse and *AC132671* in the rat) occupied a prominent position. Compared to its expression in their respective controls, this gene was the most significantly downregulated in the rat/STREP, rat/IDPN-3wk and rat/IDPN-7wk groups, and was ranked 14 in the mouse/IDPN and 75 in the rat/IDPN-4wk groups. With combined RNAscope *in situ* hybridisation and MYO7A immunohistochemistry we noticed that many, but not all, HCs expressed *Vsig10l2* (Fig 12A). By measuring fluorescence intensity, we corroborated that these HCs had a large decrease in *Vsig10l2* mRNA expression in rat/IDPN-4wk samples (Fig. 12 A,C). While the number of HCs per image was not affected by IDPN exposure neither in the centre of the crista (Fig. 12Ca, t=0.75, 6 df, p=0.38) nor in its periphery (Fig. 12Cc, t=0.06, 6 df, p=0.96), the fluorescence intensity per HC of the *Vsig10l2* mRNA was decreased in both areas (Fig 12Cb, centre: t=4.99, 6 df, p=0.002; Fig. 12Cd, periphery: t=6.91, 6 df, p=0.0004). In contrast, IDPN did not change the signal associated with the control gene *Ppib* mRNA (Fig 12Cb,d, centre and periphery p values were 0.53 and 0.72, N.S.). We also evaluated immunofluorescence associated with an anti-VSIG10L2 rabbit antiserum, in co-labelling experiments with anti-osteopontin and anti-calretinin antibodies. The results showed anti-VSIG10L2 immunoreactivity in HCIs but not in HCIIs (Fig 12B), and a strong decrease in immunoreactivity in utricle samples of IDPN-4wk rats. (Fig 12 B,D).

**Figure 12.**
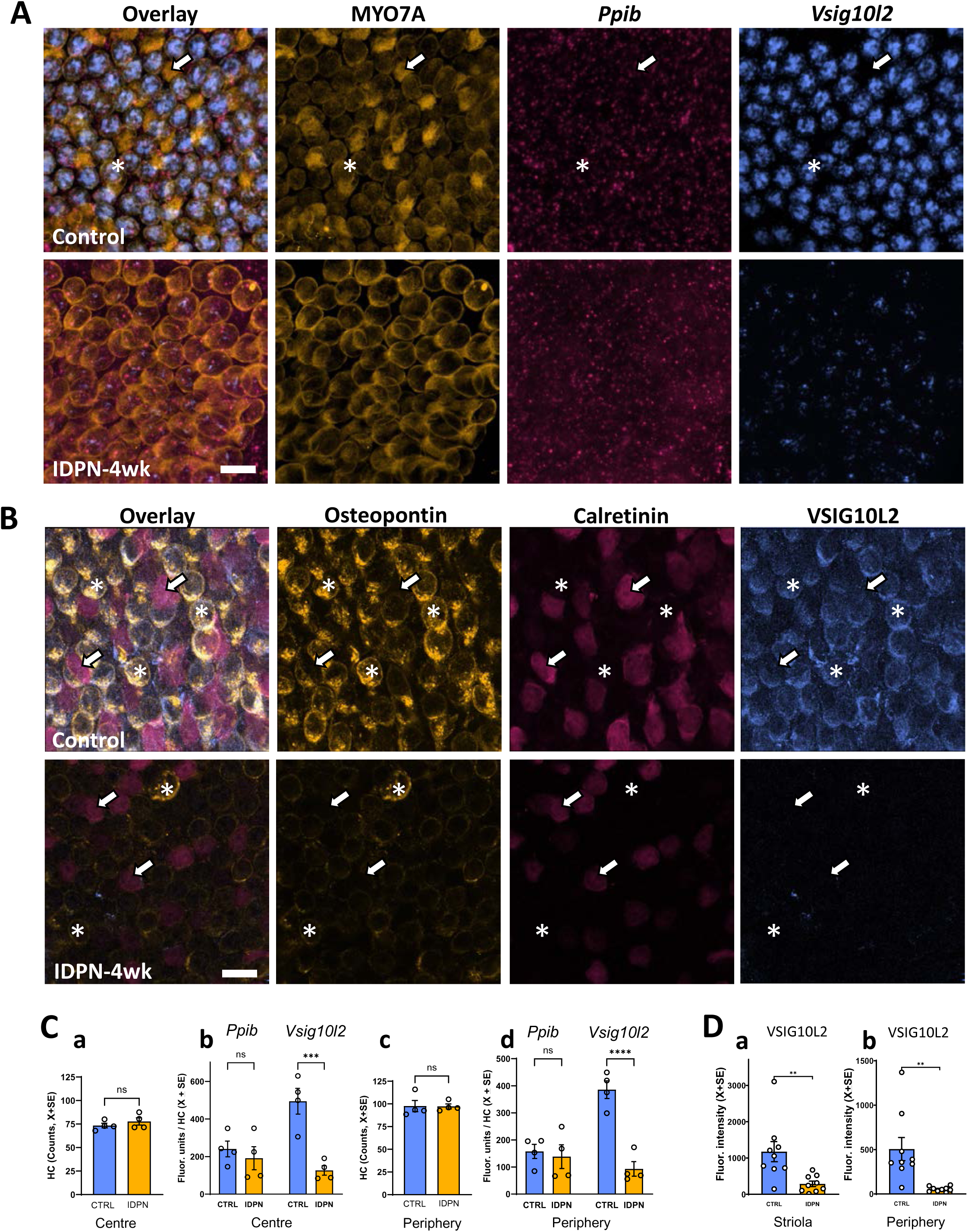
Effect of subchronic IDPN ototoxicity on *Vsig10l2* expression in HCs. **(A).** Representative images of mRNA expression analysis of *Vsig10l2* and of the control gene *Ppib* by RNA-scope, together with MYO7A immunohistochemistry in the peripheral crista of control (upper row) and IDPN-4wk (lower row) rats. The asterisks and arrows label one example of HC expressing or not *Visg10l2*, respectively. Note the loss of the *Vsig10l2* mRNA label after ototoxicity. Scale bar: 10 μm. **(B).** Representative images of osteopontin, calretinin, and VSIG10L2 immunoreactivity in the peripheral utricle of control (upper row) and IDPN-4wk (lower row) rats. Note that VSIG10L2 is expressed in HCI (osteopontin+, asterisks) but not HCII (calretinin+, arrows). Note also the decrease in label for all three antibodies. Scale bar: 10 μm. **(C).** Quantitative analysis of the number of HCs **(a,c)** and of the RNA-scope puncta **(b, d)**, comparing fluorescence units of *Ppib* and *Vsig10l2* mRNA per HC in the centre **(a, b)** and the periphery **(c, d)** of the crista of control and IDPN-4wk rats. **(D).** Quantitative analysis of anti-VSIG10L2 immunoreactivity, in the **(a)** striola and **(b)** periphery regions of the utricle of control and IDPN-4wk rats. In **C** and **D**, the bars show mean +/- SEM values and each data point is from an individual animal. **: p<0.01, ***: p<0.001, **** : p<0.0001, ns: not significant, Student’s t-test.

We also examined the effect of streptomycin on *Vsig10l2* expression after exposure *in vitro*. As shown in Fig. 13, exposure of vestibular cristae at low concentrations (0.05 μM) of streptomycin for 7 days did not cause significant loss of HCs (centre: t_6df_=0.410, p=0.70; periphery: t_6df_=0.56, p=0.59) or decreased expression of the control gene *Ppib* (centre: t_6df_=0.056, p=0.95; periphery: t_6df_=1.83, p=0.12). It did reduce the expression of the *Vsig10l2* gene in the HCs, both in the centre (t_6df_=2.92, p=0.026) and the periphery (t_6df_=6.95, p<0.001) of the crista.

**Figure 13.**
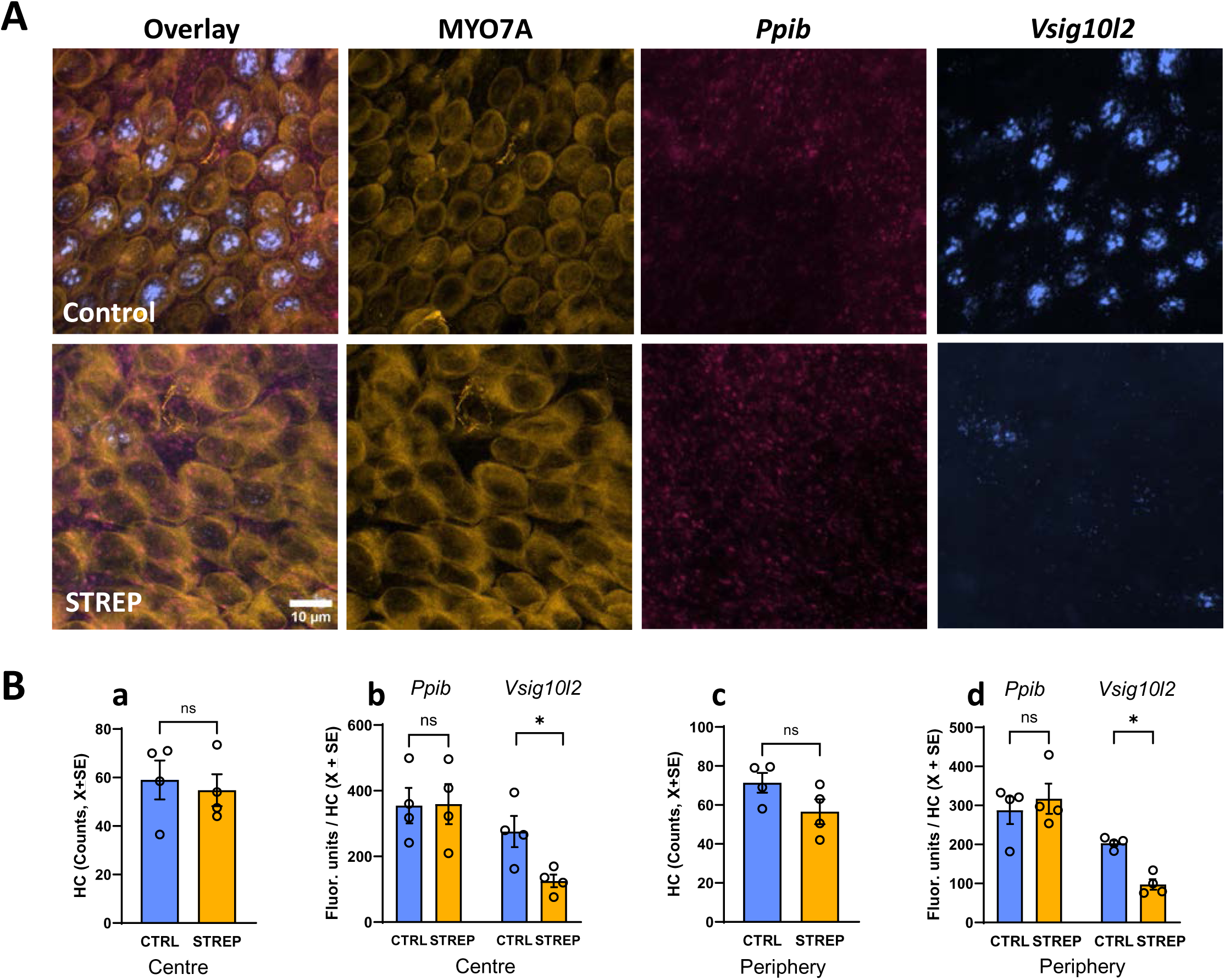
Effect of long term low dose streptomycin exposure on *Vsig10l2* expression in the rat vestibular crista *in vitro*. The cristae were placed in culture on postnatal day 1, allowed to mature for 14 days, and exposed to 0.05 μM streptomycin on days 14 to 21. **(A).** Representative images of the expression of *Vsig10l2* mRNA expression and the control gene *Ppib* by RNA-scope, together with MYO7A immunohistochemistry in the peripheral area of the control (upper row) and streptomycin (lower row) cristae. Scale bar: 10 μm. **(B).** Quantitative analysis of the number of HC in the area **(a, c)** and the puncta of RNA-scope **(b, d)**, comparing fluorescence units of the *Ppib* and *Vsig10l2* mRNA per HC in the centre **(a, b)** and the periphery **(c, d)** of the crista. The bars show mean +/- SEM values and each data point is from an individual epithelium. *: p<0.05, ns: non significant, Student’s t-test.

#### HCs under chronic stress increase ATF3 expression

RNAseq data identified *Atf3* as the first stress gene upregulated by IDPN and streptomycin in the vestibular sensory epithelium of rats and mice, so we aimed to determine whether this corresponded to a HC response. To this end, RNAscope puncta of *Atf3* and *Ppib* (control gene) were quantified in MYO7A-positive cells (Fig 14). Quantitative data (Fig14B) demonstrated that a specific increase in *Atf3* mRNA expression occurs in rat HCs after exposure to IDPN-4wk. Although the density of *Ppib* puncta per HC was not affected either in the centre (t_6df_=0.496, p=0.638) or in the periphery (t_6df_=1.388, p=0.215) of the crista, significant increases were recorded for *Atf3* expression in both areas (centre: t_6df_=3.043, p=0.0023; periphery: t_6df_= 2.659, p=0.0376. In this experiment, a modest (18%) but significant decrease in the number of HCs was recorded in the periphery (t_6df_=3.469, p=0.013) but not in the centre (t_6df_=0.080, p=0.939) of the crista.

**Figure 14.**
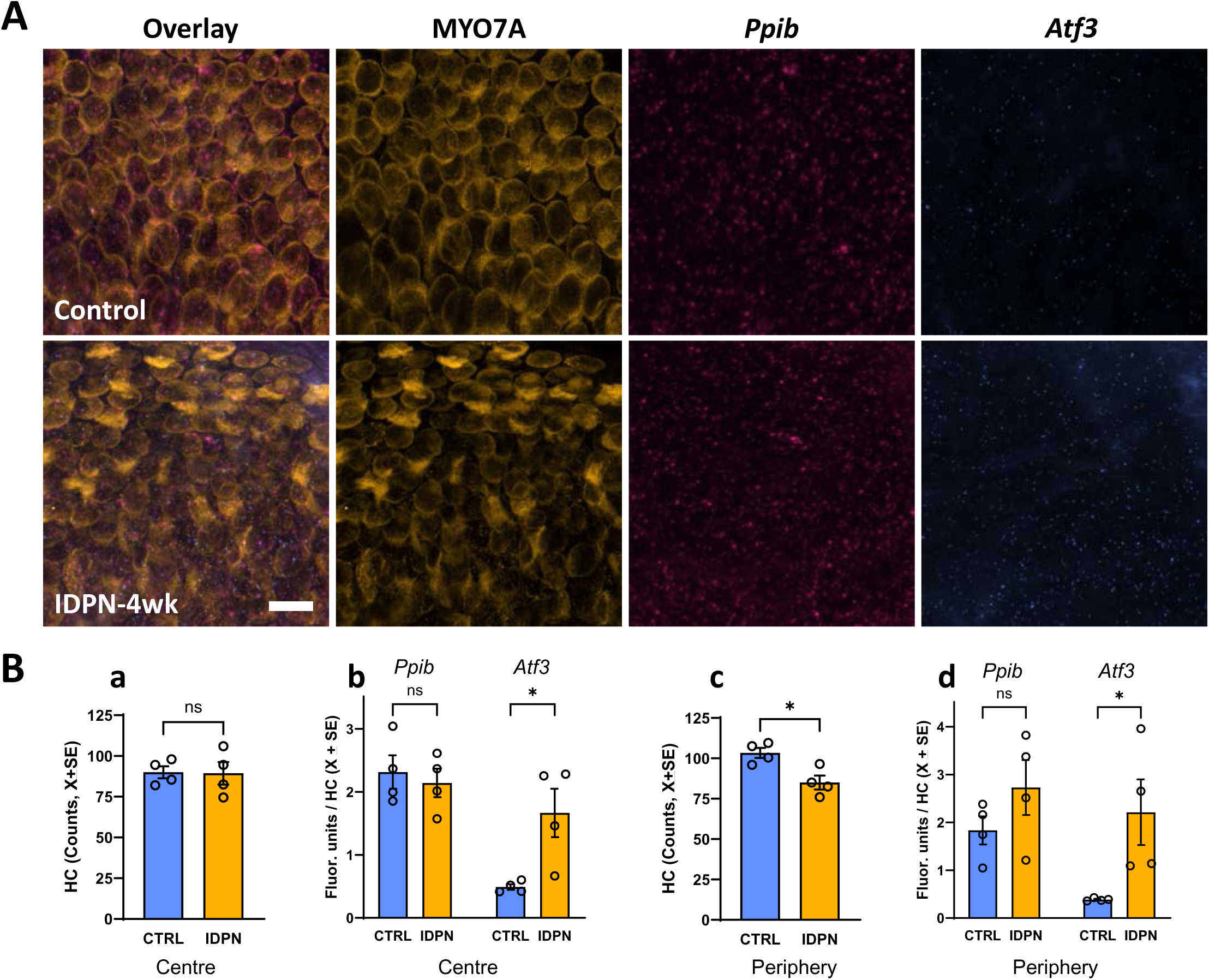
Effect of subchronic ototoxicity on *Atf3* expression in HCs. **(A).** Representative images of analysis of *Atf3* mRNA expression and the control gene *Ppib* by RNA-scope, together with MYO7A immunohistochemistry in the peripheral crista of control rats (upper row) and IDPN-4wk rats (lower row). Scale bar: 10 μm. **(B).** Quantitative analysis of the number of HCs in the area **(a, c)** and of RNA-scope puncta **(b, d)**, comparing fluorescence units of *Ppib* and *Atf3* mRNA per HC in the centre **(a, b)** and the periphery **(c, d)** of the crista of control and IDPN-4wk rats. The bars show mean +/- SEM values and each data point is from an individual animal. *: p<0.05, ns: nonsignificant, Student’s t-test.

## DISCUSSION

The molecular mechanisms involved in HC damage from ototoxic exposure have been the subject of extensive research efforts (reviewed by Kros and Steyger, 2019). However, most of these efforts have focused on the identification of events that lead to rapid degeneration of HCs, particularly apoptosis, by exposure to high doses or by using high concentrations in *in vitro* models. In the present study, our aim was to identify the main gene expression changes associated with early physiopathological response of the vestibular epithelium to sustained ototoxic stress, which ultimately leads to HC extrusion. We hypothesize that this response is more relevant to human health than those revealed by standard *in vitro* or acute high-dose exposure models. To better target the most fundamental responses, we conducted a comparative study using different species and ototoxic compounds. The data collected reveal new mechanisms that may play an important role in HC damage and loss.

An important first aspect to consider is the relative occurrence of extrusion *versus* apoptosis in the exposure models used in the study. Although there are ototoxicity models in which both extrusion and apoptosis are simultaneously observed (Li et al., 1995), exposure of male Long-Evans rats to 20 mM of IDPN in drinking water causes extrusion with no simultaneous occurrence of apoptosis (Seoane et al., 2001), even though HC apoptosis is predominant after sub-acute exposure to the same compound (Seoane et al., 2001). In support of this low incidence of apoptosis in our exposure models, the GSEA analyses (table 3) indicated that the apoptosis hallmark gene set was not upregulated in the mouse/IDPN and rat/STREP samples and modestly upregulated only in the rat/IDPN samples. Another greatly interesting aspect of the rat 20 mM IDPN model is that little to no HC loss occurs for up to 4 weeks of exposure and that the changes observed at this time point are reversible (Seoane et al., 2001; Sedó-Cabezón et al., 2015; Martins-Lopes et al., 2019). This offers the opportunity to study a population of similarly stressed cells that have not yet been extruded, as proven by the extensive loss of CASPR1 immunostaining and simultaneous deep loss of vestibular function, but with observed recovery in both parameters (Sedó-Cabezón et al., 2015). According to our previous data, the mouse/IDPN (Greguske et al., 2019) and rat/STREP (Maroto et al., 2023) models show less synchrony in the HC response, with some overlap of HC detachment and HC loss, but nevertheless show robust CASPR1 loss in association with vestibular function loss. Therefore, our working hypothesis was that the comparison of the rat/IDPN, mouse/IDPN, and rat/STREP models would inform the fundamental response of HCs to mild sustained stress. Inclusion of the rat/IDPN-3wk as an early time point was intended to reinforce the focus on the early stages of the HC response, preceding HC extrusion. In contrast, we included a time point (rat/IDPN-7wk) in which the HC extrusion is actively developing (Seoane et al., 2001; Sedó-Cabezón et al., 2015; Martins-Lopes et al., 2019) to attempt to identify gene expression responses related to extrusion.

The transcriptomic data presented here bore a batch effect that prevented the direct merging of the raw data for comparison. This came as no surprise, because the four RNA-seq experiments, encompassing five control *vs.* treated comparisons, were carried out over several years in parallel with the experiments that characterized the mouse/IDPN, rat/IDPN and rat/STREP models published elsewhere (Sedó-Cabezón et al., 2015; Greguske et al., 2019; Martins-Lopes et al., 2019; Maroto et al., 2023). However, comparisons of the DEGs and GO term lists for all experiments provided meaningful information on the response of the vestibular epithelium to chronic ototoxic stress, as summarized in Fig. 15. These comparisons revealed common gene expression changes that open new paths in our understanding of the mechanisms that lead to the loss of HCs. In addition to its interest in the biology of HCs, this knowledge may guide the identification of targets for effective treatments to prevent or block damage progression.

**Figure 15.**
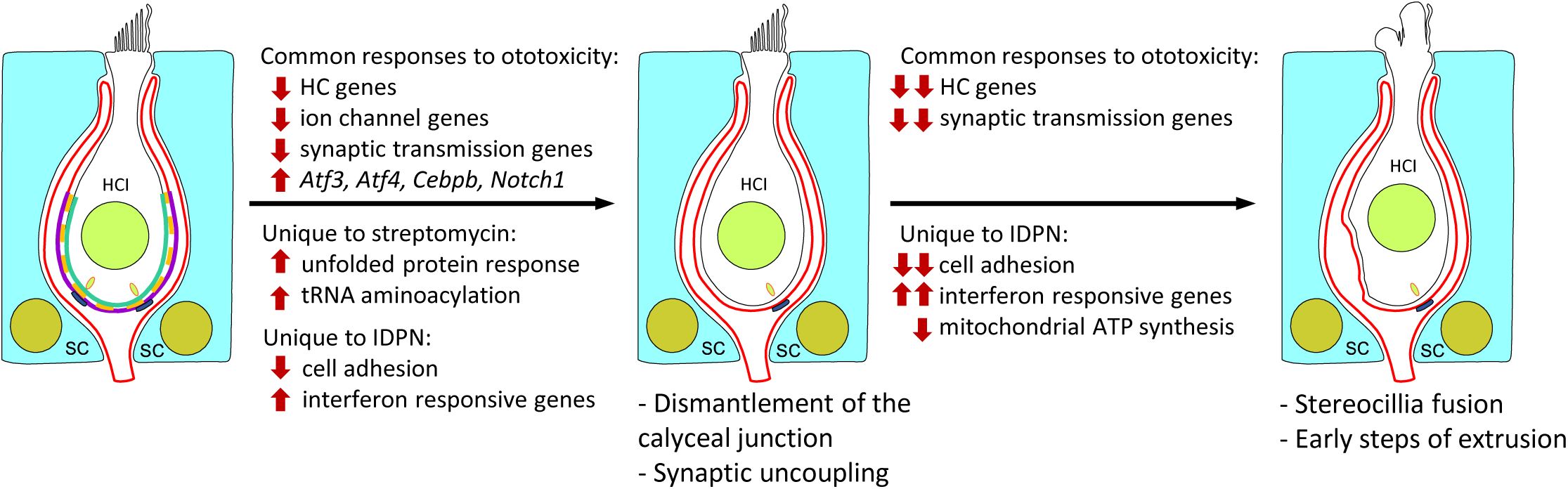
Summary of key gene expression responses identified in association with early stages of HC damage during chronic vestibular toxicity. A single HCI with two supporting cells is represented. HCII do not have calyceal junctions and are affected later during chronic ototoxicity, but they finally show similar synaptic uncoupling, ciliary coalescence, and extrusion.

Individual analyses of the transcriptomic data (Figs. 2 and 3) demonstrated that each of the exposure conditions caused a gene expression response: in each experiment control animals and treated animals were grouped by their gene expression profiles. The modest sample size (n=3/group) may have limited the ability to detect relevant subtle differences, so meaningful gene expression responses may have escaped identification. In addition, the use of bulk analysis to avoid interference of the cellular stress associated with cell isolation protocols, inevitably causes a masking effect with no meaningful differences in expression being detected in one cell type due to the dilution of the corresponding RNAs into the RNA from all cell types. Therefore, it is possible that experiments with single cell or nuclei sequencing or using larger sample sizes could have identified responses not recognized here. However, these same limitations support the robustness of the identified responses, in particular those detected in different experimental models.

Comparison across experiments revealed that the earliest and most consistent response is a robust downregulation of HC-specific genes, already present at the mildest exposure condition (rat/IDPN-3wk), and broadening in longer exposures. Because we used a bulk mRNA analysis from whole tissue, the loss of HC-specific mRNAs could be a consequence of the physical loss of HCs. However, this was clearly not the case, as the selected models offer data from stressed but still viable HCs according to previous studies (Sedó-Cabezón et al., 2015; Greguske et al., 2019; Maroto et al., 2023). The data in Fig. 1E-G also support this conclusion. The counts of hair bundles in SEM images in Fig. 1E corroborate that most HCs were present in the samples used for the RNA-seq, although data from mice permitted to recover from IDPN exposure suggest that some of these HCs were destined to extrude from the epithelium. In the rat/IDPN-4wk model, we found HC loss only in the centre of the crista (Fig. 1F). The central region of the crista is estimated to cover 1/3 of its area and has a HC density 30% less than its peripheral area. As the RNA-seq data were obtained from pooled tissue - 6 cristae and 2 utricles from individual animals-, and the total HC numbers of each type of epithelium are available (Borrajo et al., 2024), we calculated that the sequenced rat/IDPN-4wk samples had a maximum loss of HCs of 4%. In this model, no further loss is recorded if the animals were to be studied after a recovery period. Also, HC loss in the rat/IDPN-3wk samples cannot exceed the 4% calculated for the rat/IDPN-4wk. Evidently, rat/IDPN-7wk samples were likely to suffer a significant loss of HCs, but the shorter exposed groups ensure the robustness of the conclusion. In the rat/STREP samples, no statistically significant differences were obtained in HC counts, but a 10-20% loss may occur. Together, the data demonstrated a limited incidence of HC loss, which is unable to account for the observed decreases in RNA, and subsequent validation studies reinforced this conclusion.

In the gene and protein expression studies, decreases in the mRNA and protein content were demonstrated at the HC level in rat/IDPN and rat/STREP samples. Furthermore, a significant decrease in *Vsig10l2* expression was demonstrated in vestibular epithelia treated *in vitro* for 1 week with low streptomycin concentrations that do not cause HC loss. Taken together, these data firmly demonstrate that vestibular HCs under sustained stress downregulate the expression of HC-specific genes, including *Vsig10l2*, *Bdnf*, *Bmp2*, *Atp2b2*, *Dner*, *Kcna10, Xirp2,* and *Ptprq*. Although the validation data were obtained in rats only, a comparison of the gene expression data for rats and mice (Figs. 3-5) supports the conclusion that the same gene expression response occurs in mice.

In previous behavioural and histological studies of chronic ototoxicity (Sedó-Cabezón et al., 2015; Greguske et al., 2019; Maroto et al., 2023) we concluded that a reversible suspension of HC function (transduction and synaptic transmission) precedes overt signs of HC damage, such as stereocilia fusion. The transcriptomic response recorded in the vestibular ganglion (Greguske et al., 2023) also supports this phenomenon of silencing of HC. The present data indicate that this early step in chronic ototoxicity includes a downregulation of selective HC genes, including stereocilium, synaptic, and ion channel genes. This change in the gene expression phenotype of HCs under stress is congruent with the subsequent loss of specific HC features shown by the cells as they progress towards extrusion. It should be noted that this change may add difficulty in the interpretation of expression data of damaged vestibular epithelia in search of evidence of atypical HCs that could denote HC regeneration (Taylor et al. 2018; Wang et al., 2024).

Of the 34 genes downregulated under all exposure conditions, 29 have previously been identified to be selectively expressed by HCs within the rodent or human vestibular sensory epithelium (Scheffer et al., 2015; Sadler et al., 2020; Wu et al., 2013; Burns et al., 2015; Wang et al., 2024). The other 5 genes have not been identified as HC or as supporting cell specific genes, but one of them has a role in stereocilium development (*Cfap126*; Gegg et al, 2014) and another one causes hearing impairment when mutated (*Fam107b*; White et al., 2013), so it is likely that these are also HC genes. Likewise, it is tempting to speculate that the other 3 non-ascribed genes (*Ntrk3*, *Osbpl11*, *Rasal2*) are also expressed by HCs rather than SCs.

More HC-specific genes were identified in the list of 142 downregulated genes in both the rat/STREP and rat/IDPN-4-wk groups. By GO analysis, these genes are mostly associated with HC-specific BP (e.g. sensory perception of sound) and CC (e.g. stereocilium bundle) terms (Fig. 6B), strengthening the conclusion that downregulation of HC-specific genes is the main response of HCs to chronic ototoxic stress. We hypothesize that the list also contains genes with significant roles in HC differentiation or mature function that have not yet been studied. It should be noted that several of the more significantly downregulated genes have been identified as being selectively expressed in HCI versus HCII by immunohistochemistry in rodents (PMCA2, product of the *Atp2b2* gene, Borrajo et al., 2024) or by transcriptomic analysis in humans (*Adam11*, *Atp2b2*, *Bdnf*, *Bmp2*, *Chgb*, *Fbxw7*, *Kcnab1*, *Ocm*/*Ocm2*, *Vsig10l2;* Wang et al., 2024). Although genes expressed by both HCIs and HCIIs, such as *Dner,* were also identified, the data likely reflect the greater susceptibility of HCIs with respect to HCIIs to ototoxicity caused by IDPN (Maroto et al., 2021a) or streptomycin (Maroto et al., 2023). Another aspect worth discussing is the downregulation of the *Atp2b2* and *Atp2b3* genes, encoding the calcium-dependent ATP-ases, PMCA2 and SERC2. In addition to the essential role of PMCA2 in mechanotransducer function in the stereocilium (Fettiplace and Nam, 2019), reduced activity of these calcium pumps can cause cytosolic calcium overload that triggers cell death (Fettiplace and Nam, 2019; VanHouten et al., 2010).

Among the genes more robustly downregulated by chronic ototoxicity, we identified a gene, *Vsig10l2* (V-set and immunoglobulin domain containing 10 like 2), for which there was virtually no information. Automatic annotation in UniProt predicts a signal peptide and Alpha-fold (Jumper et al., 2021; Varadi et al., 2024) predicts a cell-adhesion protein with five immunoglobulin domains, one fibronectin-type III domain, and one transmembrane domain. The RNAscope and immunohistochemical data shown in Figs. 12 and 13 demonstrated that this gene is selectively expressed in HCIs and is strongly downregulated after ototoxic stress. While the role of this gene in HCI function remains to be elucidated, its interest as a marker of HCI stress becomes obvious. Recent studies in human vestibular epithelium have also identified the *Vsig10l2* gene as a specific HCI marker (Wang et al., 2024).

Downregulation of HC-specific genes preceded a second response identified in both rats and mice and after either exposure to streptomycin or IDPN: increased expression of *Atf3*, a well-known hub mediator of cell stress responses (Heinrich et al., 2024) and related genes. Surprisingly, only recently has ATF3 been identified, using HEI-OC1 cells, as a candidate player in aminoglycoside (Zhang et al., 2022) and cisplatin (Lee et al., 2022) ototoxicity. More recently, increased *Atf3* expression has been recorded 1 h after noise exposure in cochlear tissue and in HEI-OC1 cells after stress triggered by tert-butyl hydroperoxide exposure (Liang et al., 2024). The work of Zhang et al. (2022) demonstrated increased levels of *Atf3* mRNA and protein in cells exposed to neomycin and presented evidence that this increase could affect mitophagy by transcriptional inhibition of the mitophagy regulator *Pink1*. However, we did not record changes in *Pink1* expression in the present experiments. As a stress mediator, *Atf3* promotes survival/repair or cell death responses (Hunt et al., 2012), so more research is needed to determine the potential role of this transcription factor in driving the reversible detachment and synaptic uncoupling shown by HCs in the early stages of chronic toxicity or in their subsequent determination to elimination by extrusion. In any case, the absence of upregulation of *Atf3* in the rat/IDPN-3wk samples makes it unlikely that it plays a role in early downregulation of HC-specific genes identified here.

In addition to common responses, our data also revealed some responses unique to each ototoxicity model. When discussing these results, we must keep in mind that the present work does not include experiments determining whether these effects were associated with HCs or other types of cells in the epithelium used for RNA extraction. Beyond this cautionary note, one first conclusion is that the existence of these model-specific responses reinforces the value of the responses previously identified in both models as fundamental responses of the HC to different kinds of toxic stress. By contrast, the unique responses of each model can offer clues to compound-specific toxicity mechanisms. This may be the case for the upregulation of the unfolded protein response and tRNA aminoacylation BPs after streptomycin, which may possibly be related to the effect of aminoglycosides on ribosomal translation (O’Sullivan et al., 2017). Similarly downregulation of mitochondrial ATP synthesis might point to a nitrile-specific effect. However, a number of facts suggest that the two robust and time-increasing responses found after IDPN exposure and not observed in the streptomycin model may be of broad, rather than IDPN-specific, significance. First, downregulation of cell adhesion genes fits well with the loss of cell adhesion contributing to HC extrusion, a phenomenon observed not only after chronic IDPN (Seoane et al., 2021a,b; Sedó-Cabezón et al., 2015; Greguske et al., 2019), but also after chronic streptomycin (Maroto et al., 2023). In these studies, a marked contrast was observed between the slow and highly synchronous process of HC extrusion triggered by IDPN and the likely fast and asynchronous extrusion of HC after streptomycin (Maroto et al., 2023). Thus, it is possible that streptomycin also triggers downregulation of cell adhesion genes but that not enough affected cells are simultaneously present in the epithelium for the detection of this effect in bulk analysis. The second important dissimilar response is the upregulation of interferon responsive genes in an anti-viral-like or inflammatory-like response. Increased expression of interferon-responsible genes can occur following toxic stress, through nitrosylation of ISG15 and subsequent activation of the ISGylation process (Zhang and Zhang, 2011). Alternatively, such a response can also be triggered by STAT1 phosphorylation induced by oxidative stress, and, in fact, this has been reported to occur in HCs after exposure to cisplatin or aminoglycoside (Schmitt et al., 2009; Levano and Bodmer, 2015; Kaur et al., 2016; Borse et al., 2017). Again, the bulk epithelium approach used here, selected to circumvent the confounding effects of stress caused by cell isolation protocols, may have lacked the sensitivity to detect this response in streptomycin samples. Although still speculative, the hypothesis that decreased expression of cell adhesion genes and increased expression of interferon-regulated genes are responses of broad significance in chronic vestibular dysfunction is supported by the recent work by Kulasooriya et al. (2025). Using single-cell transcriptomics to compare young and old vestibular HCs in the mouse, these authors identified in the old samples, among other responses, 1) downregulation of hair bundle, mechanotransduction, synapse, and HC-specific genes; 2) downregulation of cell-cell adhesion genes; and 3) increase in pro-inflammatory genes indicating potential maintenance of a prolonged low-grade inflammatory state. Another study (Paplou et al., 2023) has also reported decreased expression of cell adhesion genes and increased expression of inflammation-related genes, although the use of the entire vestibule (bone-encapsulated) likely reduced the sensitivity of the study. Therefore, these may be common responses of HCs to chronic stress. In support of this conclusion, we have recorded cellular evidence of decreased HC adhesion in human vestibular epithelium affected by a neighbouring tumour likely causing inflammatory stress (Maroto et al., 2023).

In summary, the present study compared the RNA-seq results of four experiments involving five control *vs.* chronic ototoxicity conditions. The data obtained identified that marked downregulation of HC-specific genes occurs as the first response to chronic stress. Among well-known stress responses, upregulation of *Atf3* was recorded in both rats and mice and after exposure to IDPN and streptomycin. In contrast, other prominent responses were identified after one compound-specific exposure, notably the unfolded protein response identified in streptomycin exposed rats, and the down regulation of cell adhesion and upregulation of an anti-viral-like response identified in rats and mice exposed to IDPN. These require further research to establish their link to particular mechanisms of ototoxicity or reveal their broader role. This study opens new paths in our understanding of the mechanisms that lead to the loss of HC during chronic stress conditions.

## Acknowledgements

M.B, E.A.G, A.F.M, A.P., A.B.-G., and J.L. were members of the SGR Group 2017SGR621 (Agència de Gestió d’Ajuts Universitaris i de Recerca, AGAUR, Generalitat de Catalunya). The confocal microscopy studies were performed at the Centres Científics i Tecnològics de la Universitat de Barcelona (CCiTUB). We thank Dr. Benjamín Torrejon-Escribano and Dr. Ashraf Muhaisen for their expert and technical help, and students Clara López Escolà, Javier Domínguez Rovira, Júlia Valor Blanquer and Maria Capdevila Bayo for their contributions towards the studies. We also thank Dr. Anna Lysakowsi for advice on KCNA10 labelling.

## Ethics approval

The use of animals followed the local regulations (Law 5/1995 and Act 214/1997 of the Generalitat de Catalunya, and Law 6/2013 and Act 53/2013 of the Gobierno de España) in agreement with the EU Directive 2010/63. The experimental protocols were approved by the Universitat de Barcelona’s Ethics Committee on Animal Experiments and the Commission on Animal Experimentation of the Generalitat de Catalunya (numbers 10913 and 10328). No human subjects or samples were involved in the study.

