## Supplementary Figure S1 for "Early downregulation of hair cell (HC)-specific genes in the vestibular sensory epithelium during chronic ototoxicity"

Suppl. Fig S1.

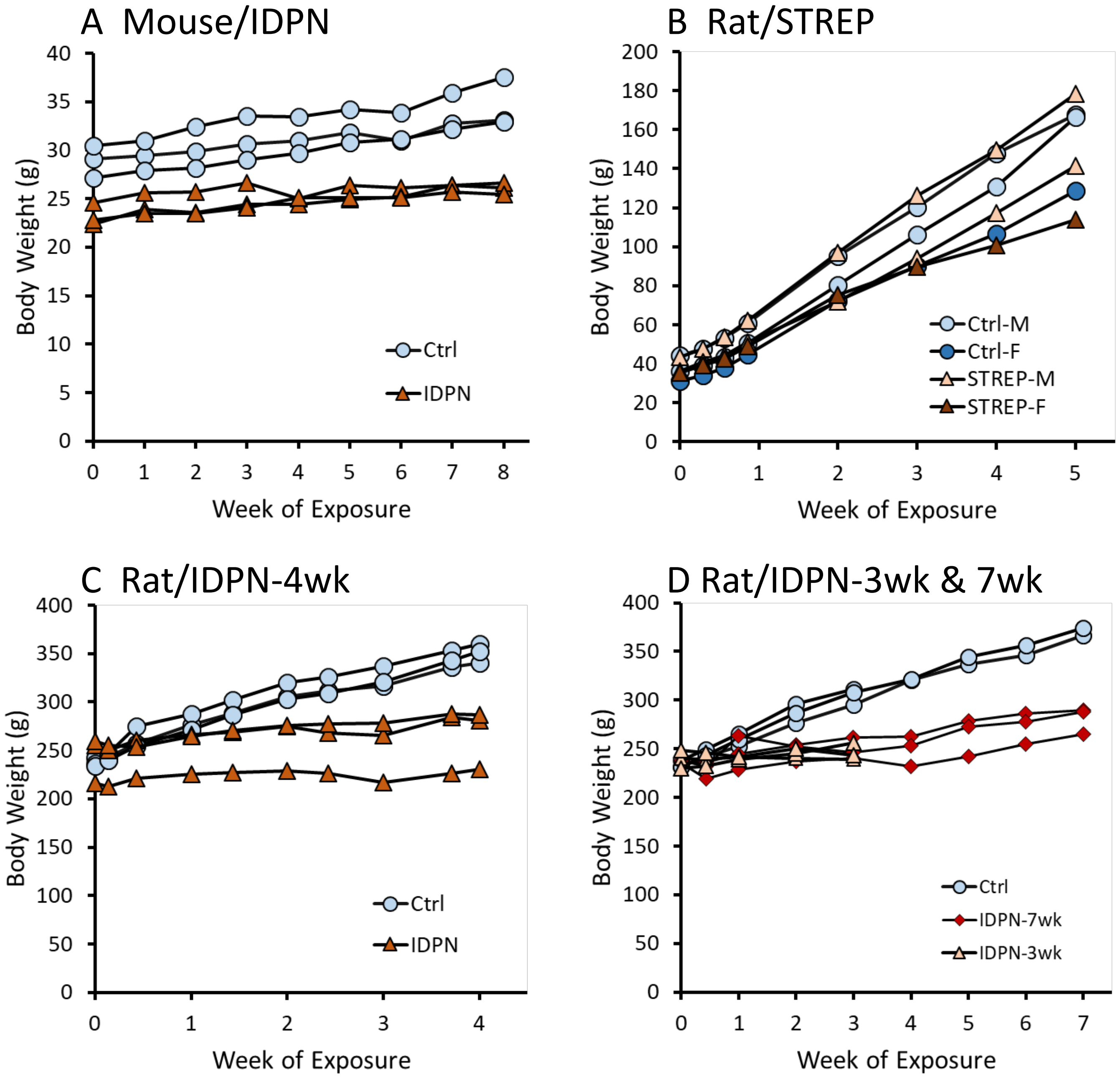

A Mouse/IDPN
Body Weight (g)
Week of Exposure
Ctrl
IDPN
B Rat/STREP
Body Weight (g)
Week of Exposure
Ctrl-M
Ctrl-F
STREP-M
STREP-F
C Rat/IDPN-4wk
Body Weight (g)
Week of Exposure
Ctrl
IDPN
D Rat/IDPN-3wk & 7wk
Body Weight (g)
Week of Exposure
Ctrl
IDPN-7wk
IDPN-3wk

Supplementary Fig S1. Effect of subchronic otototoxicity on body weight. Graphs show the body weight of the individual animals from which the RNA was extracted for RNA-seq analysis.
